# Loss of TREM2 reduces hyperactivation of progranulin deficient microglia but not lysosomal pathology

**DOI:** 10.1101/2021.07.08.451574

**Authors:** Anika Reifschneider, Sophie Robinson, Bettina van Lengerich, Johannes Gnörich, Todd Logan, Steffanie Heindl, Miriam A Vogt, Endy Weidinger, Lina Riedl, Karin Wind, Artem Zatcepin, Sophie Haberl, Brigitte Nuscher, Gernot Kleinberger, Julien Klimmt, Julia K Götzl, Arthur Liesz, Katharina Bürger, Matthias Brendel, Johannes Levin, Janine Diehl-Schmid, Jung Suh, Gilbert Di Paolo, Joseph W Lewcock, Kathryn M Monroe, Dominik Paquet, Anja Capell, Christian Haass

## Abstract

*GRN* haploinsufficiency causes frontotemporal lobar degeneration and results in microglial hyperactivation, lysosomal dysfunction and TDP-43 deposition. To understand the contribution of microglial hyperactivation to pathology we evaluated genetic and pharmacological approaches suppressing TREM2 dependent transition of microglia from a homeostatic to a disease associated state. *Trem2* deficiency in *Grn* KO mice led to a reduction of microglia activation. To explore antibody-mediated pharmacological modulation of TREM2-dependent microglial states, we identified antagonistic TREM2 antibodies. Treatment of macrophages from *GRN*-FTLD patients with these antibodies allowed a complete rescue of elevated levels of TREM2 together with increased shedding and reduction of TREM2 signaling. Furthermore, antibody-treated PGRN deficient hiMGL showed dampened microglial hyperactivation, reduced TREM2 signaling and phagocytic activity, however, lack of rescue of lysosomal dysfunction. Similarly, lysosomal dysfunction, lipid dysregulation and glucose hypometabolism of *Grn* KO mice were not rescued by TREM2 ablation. Furthermore, NfL, a biomarker for neurodegeneration, was elevated in the *Grn*/*Trem2* KO. These findings suggest that microglia hyperactivation is not necessarily contributing to neurotoxicity, and instead demonstrates that TREM2 exhibits neuroprotective potential in this model.

## Introduction

Neurodegenerative diseases are currently incurable and novel therapeutic strategies are desperately required. Besides disease defining protein deposits (Aguzzi and Haass, 2003), microgliosis is observed in almost all neurodegenerative diseases (Ransohoff, 2016). Microgliosis can be detrimental (Heneka et al., 2013; Hong et al., 2016a). However, recent findings strongly suggested that certain microglial responses to brain pathology may also be protective (Deczkowska et al., 2020; Lewcock et al., 2020). This is based on the identification of variants in genes predominantly or exclusively expressed in microglia within the brain that increase the risk for late onset Alzheimer’s disease (LOAD) and other neurodegenerative disorders (Efthymiou and Goate, 2017). Protective microglial functions became particularly evident upon functional investigations of coding variants found within the triggering receptor expressed on myeloid cells 2 (TREM2) gene, which can increase the risk for LOAD and other neurogenerative disorders including frontotemporal dementia-like syndromes (Guerreiro et al., 2013; Jonsson et al., 2013). These TREM2 variants reduce lipid ligand binding, lipid and energy metabolism, chemotaxis, survival/proliferation, phagocytosis of cellular debris and potentially other essential microglial functions (Deczkowska et al., 2020; Lewcock et al., 2020). Moreover, a loss of TREM2 function locks microglia in a homeostatic state (Keren-Shaul et al., 2017; Krasemann et al., 2017; Mazaheri et al., 2017; Nugent et al., 2020), in which they are unable to respond to pathological challenges by inducing a disease associated mRNA signature.

Disease associated microglia (DAM) respond to amyloid pathology by clustering around amyloid plaques where they exhibit a protective function by encapsulating the protein deposits via a barrier function (Yuan et al., 2016) that promotes amyloid plaque compaction (Meilandt et al., 2020; Ulrich et al., 2014; Wang et al., 2016), and reduce *de novo* seeding of amyloid plaques (Parhizkar et al., 2019). TREM2 is therefore believed to be a central target for therapeutic modulation of microglial functions (Deczkowska et al., 2020; Lewcock et al., 2020). A number of agonistic anti-TREM2 antibodies were recently developed (Cheng et al., 2018; Cignarella et al., 2020; Ellwanger et al., 2021; Fassler et al., 2021; Price et al., 2020; Schlepckow et al., 2020; Wang et al., 2020), which either enhance cell-surface levels of signaling competent TREM2 by blocking TREM2 shedding and/or crosslink TREM2 receptors to stimulate downstream signaling via Syk phosphorylation. In preclinical studies these antibodies boost protective functions of microglia as shown by enhanced amyloid ß-peptide (A ß) and myelin clearance, reduced amyloid plaque load, improved memory in models of amyloidosis, and support of axon regeneration and remyelination in models of demyelinating disorders such as multiple sclerosis (Bosch-Queralt et al., 2021; Cheng et al., 2018; Cignarella et al., 2020; Ellwanger et al., 2021; Fassler et al., 2021; Lewcock et al., 2020; Price et al., 2020; Schlepckow et al., 2020; Wang et al., 2020).

Although, increased TREM2 may be protective in AD patients (Ewers et al., 2019), in other neurodegenerative diseases microglia may be overactivated and become dysfunctional (Heneka et al., 2013; Hong et al., 2016a; Ransohoff, 2016). Therefore, in these contexts antagonistic TREM2 antibodies may display therapeutic benefit through dampening microglial hyperactivation. A typical example for a neurodegenerative disorder where microglia are severely hyperactivated is *GRN*-associated frontotemporal lobar degeneration (*GRN*-FTLD) with TDP-43 (transactive response DNA-binding protein 43 kDa) deposition caused by progranulin (PGRN) deficiency (Baker et al., 2006; Cruts et al., 2006; Gotzl et al., 2019). In models for *GRN*-FTLD associated haploinsufficiency, hyperactivation of microglia is evident, as demonstrated by an increased disease associated mRNA signature as well as strongly increased 18kDa translocator protein positron-emission-tomography (TSPO)-PET signals in mouse models (Gotzl et al., 2019; Lui et al., 2016; Zhang et al., 2020). This is the opposite phenotype of *Trem2* knockout (KO) microglia, which are locked in a homeostatic state (Gotzl et al., 2019; Keren-Shaul et al., 2017; Kleinberger et al., 2017; Krasemann et al., 2017; Mazaheri et al., 2017; Nugent et al., 2020). Hyperactivation of microglia is also observed in the brain of *GRN*-FTLD patients (Gotzl et al., 2019; Lui et al., 2016; Woollacott et al., 2018). PGRN is a secreted protein, which is also transported to lysosomes (Hu et al., 2010; Zhou et al., 2015), where it appears to control activity of hydrolases, such as cathepsins and glucocerebrosidase (GCase) (Arrant et al., 2019; Beel et al., 2017; Gotzl et al., 2018; Gotzl et al., 2016; Logan et al., 2021; Paushter et al., 2018; Ward et al., 2017). Total loss of PGRN results in a lysosomal storage disorder (Almeida et al., 2016; Gotzl et al., 2014; Logan et al., 2021; Smith et al., 2012). A potential synergistic contribution of lysosomal dysfunction and hyperactivated microglia to the disease pathology and specifically to the deposition of TDP-43 in neurons is likely but currently not understood (Logan et al., 2021).

To determine whether hyperactivation of microglia and its pathological consequences in *Grn* KO mice are dependent on aberrant TREM2 signaling, we sought to reduce the microglial activation status by crossing them to *Trem2* KO mice. This reduced the expression of DAM genes, suggesting that negative modulation of TREM2 signaling may be exploited to lower microglial activation in neuroinflammatory disorders. In analogy to the agonistic 4D9 TREM2 antibody developed earlier in our laboratory (Schlepckow et al., 2020), we therefore generated monoclonal antibodies with opposite, namely antagonistic properties. Such antibodies blocked lipid ligand-induced TREM2 signaling, reduced signaling competent cell-surface TREM2 in *GRN*-FTLD patient derived macrophages and concomitantly increased shedding of TREM2, which resulted in enhanced release of soluble TREM2 (sTREM2). In genetically engineered human induced pluripotent stem cell-derived (iPSC) microglia lacking PGRN, the antagonistic antibodies reduced expression of the majority of candidate genes of the DAM signature but failed to restore lysosomal function. Similarly, in *Grn/Trem2* double knockout (*Double* KO) mice lysosomal dysfunction was not rescued. Moreover, pathological features such as reduced 2-deoxy-2-[18F]fluoro-d-glucose (FDG) uptake, disturbed lipid metabolism and abnormal microglial morphology were not ameliorated. Strikingly, neurofilament light chain (NfL), a sensitive fluid biomarker for neurodegeneration (Preische et al., 2019), was not reduced but even dramatically increased in the CSF. These findings therefore suggest that against common expectations, hyperactivated microglia may retain at least some TREM2 dependent protective activities.

## Results

### *Trem2* KO dampens hyperactivation of microglia in PGRN deficient mice

PGRN and TREM2 deficiency results in opposite microglial activation states (Gotzl et al., 2019). To prove if reduction of TREM2 signaling can ameliorate hyperactivation of PGRN deficient microglia, we crossed *Grn* KO mice (Kayasuga et al., 2007) to *Trem2* KO mice (Turnbull et al., 2006) and performed TSPO-PET imaging using established protocols (Kleinberger et al., 2017; Liu et al., 2015). In line with our earlier findings (Gotzl et al., 2019), we confirmed a strong increase of the TSPO-PET signal in the brain of *Grn* KO mice when compared to WT (p < 0.01) (Fig 1A and B). We also confirmed reduced TSPO expression in the brain of *Trem2* KO mice (p < 0.03) (Fig 1A and B), fitting to our findings in TREM2 loss-of-function models (Gotzl et al., 2019; Kleinberger et al., 2017). Consistent with the above-described goal to dampen hyperactivation of microglia, investigation of *Double* KO mice (Fig 1A and B) indicated a balanced TSPO expression without a significant difference when compared to WT (p = 0.945) and a reduction of TSPO expression when compared to *Grn* KO mice (p < 0.05) (Fig 1A and B).

**Figure 1.**
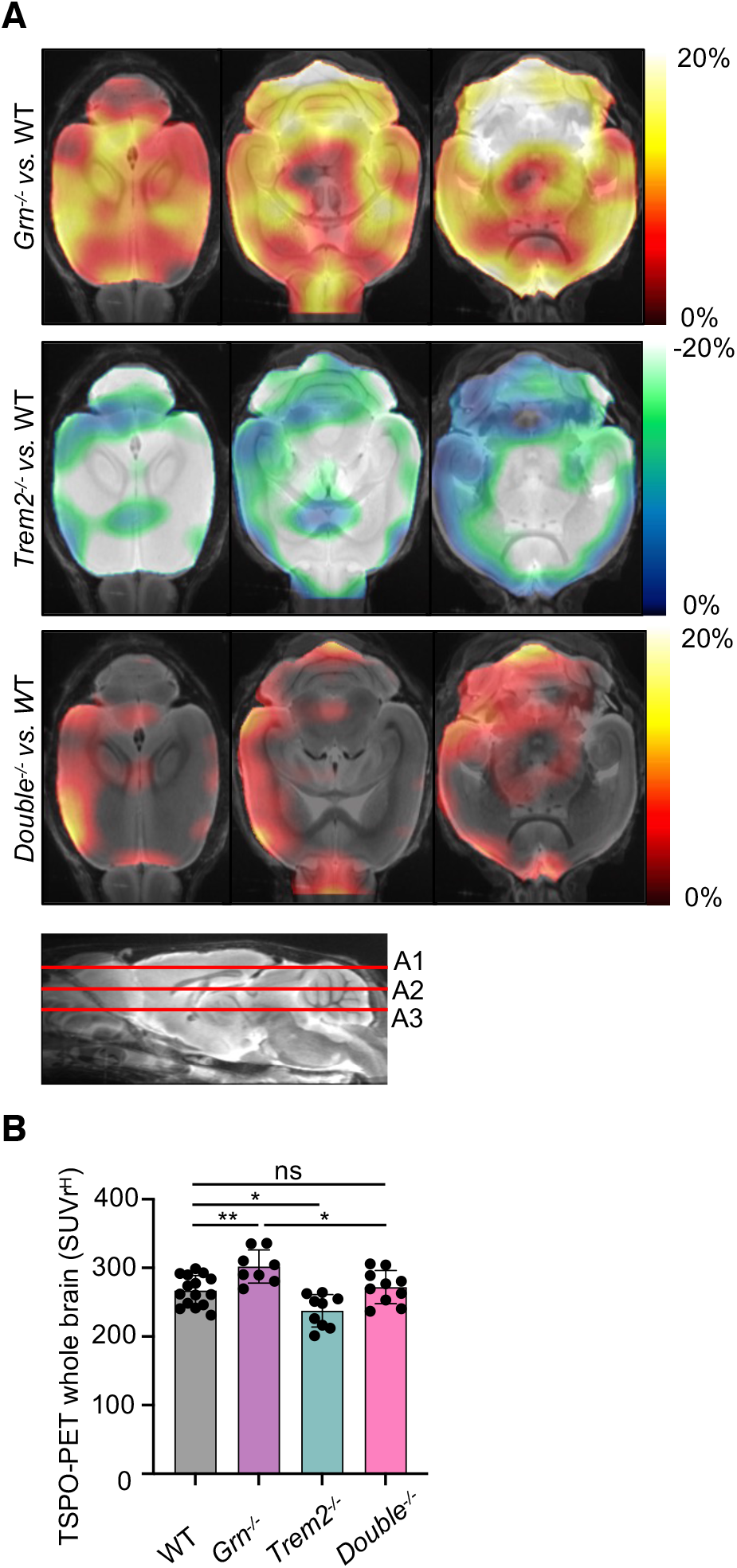
TSPO-PET imaging indicates rescue of microglial hyperactivation in *Double*^-/-^ mice. **A.** Axial slices as indicated show %-TSPO-PET differences between *Grn*^-/-^, *Trem2*^-/-^ or *Double*^-/-^ mice and WT at the group level. Images adjusted to an MRI template indicate increased microglial activity in the brain of *Grn*^-/-^ mice (hot color scale), compensated microglial activity in the brain of *Double*^-/-^ mice and decreased microglial activity in the brain of *Trem2* KO mice (cold color scale), each in contrast against age-matched WT mice. **B.** Scatter plot illustrates individual mouse TSPO-PET values derived from a whole brain volume of interest. 8-15 female mice per group at an average age of 11.1 ± 1.6 months (*Grn*^-/-^ (n = 8), *Trem2*^-/-^ (n = 9), *Double*^-/-^ (n = 10), WT (n= 15)). Data represent mean ± SD. For statistical analysis one-way ANOVA with Tukey post hoc test was used. Statistical significance was set at *, *p* < 0.05; **, *p* < 0.01; and ***, *p* < 0.001; and ****, *p* < 0.0001; ns, not significant.

The above-described findings suggest that DAM gene expression patterns as observed in *Grn* KO mice may be partially rescued in *Double* KO mice. To provide direct evidence that in *Double* KO mice the molecular signature of microglia is shifted from a DAM state towards a homeostatic state, we isolated microglia from adult mouse brains. Microglial mRNA of all three mouse lines was analyzed using a customized nCounter panel (NanoString Technologies), which includes 65 genes that previously showed opposite expression levels in *Grn* and *Trem2* KO mice (Gotzl et al., 2019; Mazaheri et al., 2017). Gene expression levels were normalized against the geometric mean of four housekeeping genes, including *Asb10*, *Cltc*, *Hprt1* and *Tubb5.* In accordance with our previous findings (Gotzl et al., 2019), candidate genes of the DAM signature such as *ApoE*, *Cd22*, *Ly9, Clec7a*, *Spp1*, and *Olfr110* were massively upregulated in *Grn* KO microglia while upregulation of these genes was suppressed in the *Trem2* KO microglia (Fig 2A-E). In the *Double* KO microglia expression of the DAM signature genes *Clec7a*, *Spp1* and *Olfr110* are fully rescued and others, such as *ApoE*, *Cd22* and *Ly9* are least partially reduced compared to the *Grn* KO (Fig 2D and E).

**Figure 2.**
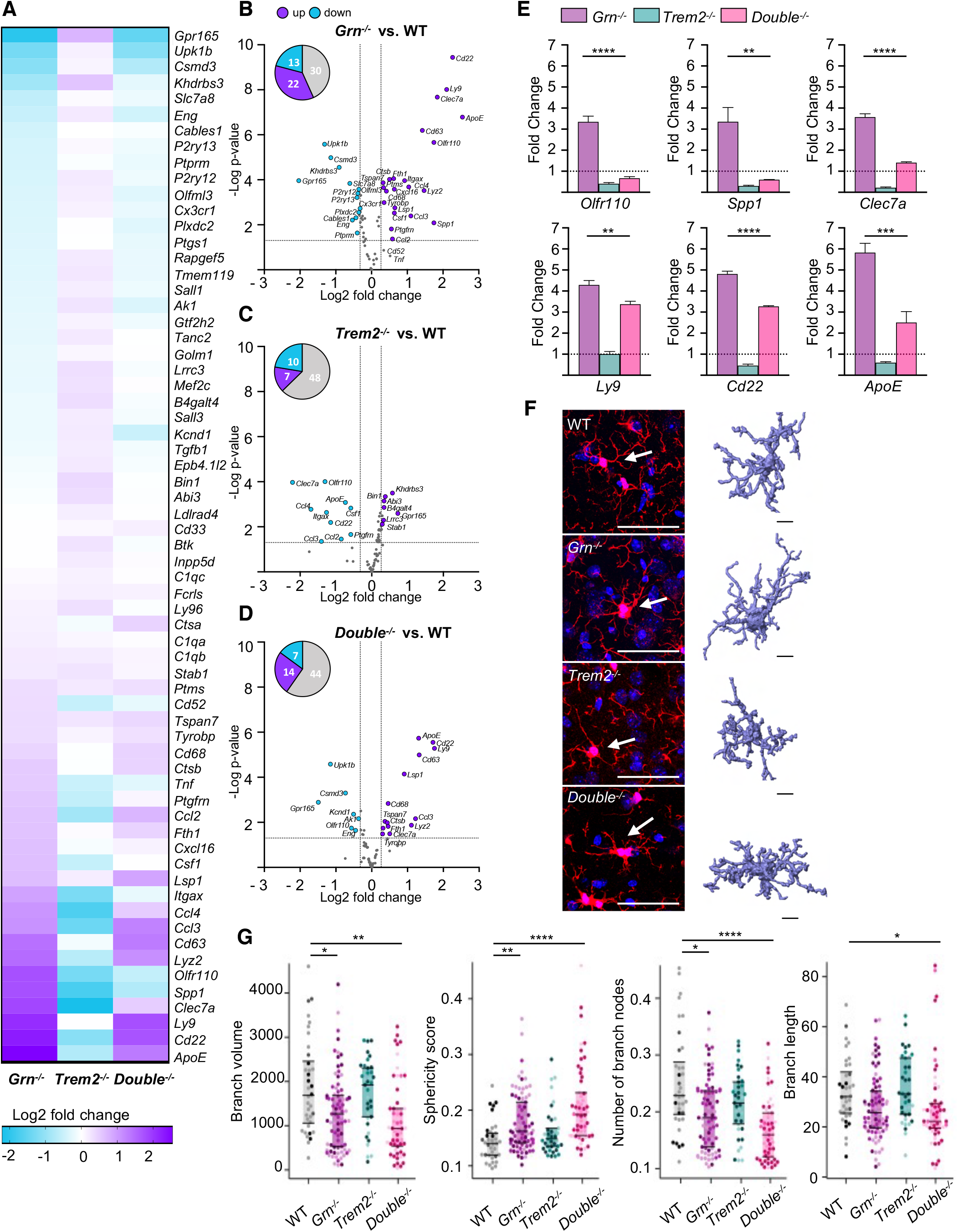
Loss of TREM2 reduces the DAM signature of *Grn*^-/-^ mice. **A.** Heatmap of 65 DAM-associated gene transcripts analyzed by NanoString in FCRLS- and CD11b-positive *Grn^-/-^* (n=6)*, Trem2^-/-^* (n=5) and *Double^-/-^* (n=6) microglia in comparison to WT (n=7) microglia isolated from 6-month-old mice. The RNA counts for each gene and sample were normalized to the mean value of WT followed by a log2 transformation. **B.** Volcano blot presentation of the differently expressed transcripts in FCRLS- and CD11b-positive *Grn^-/-^* in comparison to WT microglia isolated from 6-month-old mice. 35 out of 65 analyzed genes are significantly changed more than 20%, with 22 genes upregulated (purple) and 13 genes downregulated (blue). **C.** Volcano blot presentation of the differently expressed transcripts in FCRLS- and CD11b-positive *Trem2^-/-^* in comparison to WT microglia isolated from 6-month-old mice. 17 out of 65 analyzed genes are significantly changed more than 20%, with 7 genes upregulated (purple) and 10 genes downregulated (blue). **D.** Volcano blot presentation of the differently expressed transcripts in FCRLS- and CD11b-positive *Double^-/-^* in comparison to WT microglia isolated from 6-month-old mice. 21 out of 65 analyzed genes are significantly changed more than 20%, with 14 genes upregulated (purple) and 7 genes downregulated (blue). **E.** Expression profiles of selected DAM genes, whose mRNA levels are rescued in *Double^-/-^* vs. *Grn^-/-^,* microglia. mRNA expression normalized to the mean of the WT cohort. **F.** Morphological analysis of microglia. Representative maximum intensity projections of confocal z-stack images showing Iba1^+^ microglial cells of WT, *Grn^-/-^, Trem2^-/-^* and *Double^-/-^* mice (scalebar = 50 µm). Arrows point to individual microglia, which are shown as three-dimensional reconstruction, scalebar = 10 µm. **G.** Morphological differences of microglia from WT, *Grn^-/-^*, *Trem2^-/-^* and *Double^-/-^* mice shown by branch volume, sphericity score, branch length and the number of branch nodes. Statistical analysis of group difference for the morphological scores “Branch volume” (auc = 0.72), “Sphericity score” (auc = 0.82), “Branch length” (auc = 0.69) and “Number of branch nodes” (auc = 0.80) was performed using the Wilcoxon rank sum test with continuity correction and Bonferroni post-hoc correction for multiple testing in R (version 4.0.3). Data represent mean ± SEM. For statistical analysis in **B** - **D** the unpaired, two-tailed student’s t-test was performed, and in **E** one-way ANOVA with Dunnett’s post hoc test was used to compare *Grn^-/-^, Trem2^-/-^* and *Double^-/-^* microglia. Statistical significance was set at *, *p* < 0.05; **, *p* < 0.01; and ***, *p* < 0.001; and ****, *p* < 0.000; ns, not significant.

To obtain additional information on the activation status of *Double* KO microglia, we determined microglial morphology. We extracted morphological features in WT, *Grn* KO, *Trem2* KO and *Double* KO animals after 3D-reconstruction of Iba1^+^ microglia from confocal z-stack images (Fig 2F and G). Microglial cells from *Grn* KO and *Double* KO animals showed a significantly decreased score for “branch volume”, “number of branch nodes” and less pronounced for “branch length”, as well as a significantly increased score for “sphericity”, which is associated with an increased activation state of microglia (Heindl et al., 2018). In contrast, the morphological scores for *Trem2* KO animals were comparable to WT. Thus, although the molecular signature of hyperactivated PGRN deficient microglia is partially rescued by the loss of TREM2 function, the morphological analysis indicates that *Double* KO microglia do not rescue *Grn* KO microglial morphology.

### Antagonistic TREM2 antibodies decrease cell surface TREM2 and reduce ligand induced Syk signaling in monocyte-derived patient macrophages

Antagonist TREM2 antibodies were generated by immunizing rodents with human TREM2 extracellular domain (ECD)-Fc fusion protein and performing single B-cell sequencing on peripheral lymphoid tissues. Antibodies that bound specifically to human TREM2 were evaluated for functional impact to TREM2 signaling. Antagonistic antibodies were identified by their ability to block TREM2-dependent lipid ligand-induced activation of p-Syk on HEK293 cells over-expressing TREM2/DAP12 (Fig EV1). Cells were dosed with three different concentrations of liposomes, and antagonistic antibody 1 (Ab1) and antagonistic antibody 2 (Ab2), which were found to block PS-induced p-Syk activity (Fig EV1A). Both antibodies bind to an epitope between amino acids 29 and 63 of TREM2 (Fig 3A, Fig EV1B). These selected antibodies were reformatted onto an effectorless human hIgG1-LALAPG backbone, and demonstrated high affinity for cell surface TREM2 (0.38 nM EC50 Ab1, and 0.18 nM EC50 Ab2 in cell binding) and high affinity to human TREM2 ECD protein binding via Biacore (0.21 nM Ab1 and 4.5 nM Ab2) (Fig EV1C-G). Ligand blocking activity was further validated in human monocyte-derived macrophages, which were treated in a dose-response format with antibodies in the presence of liposomes to determine the potency of Ab1 and Ab2 to block liposome-induced TREM2-mediated p-Syk signaling (Fig 3B, Fig EV1C).

**Figure 3.**
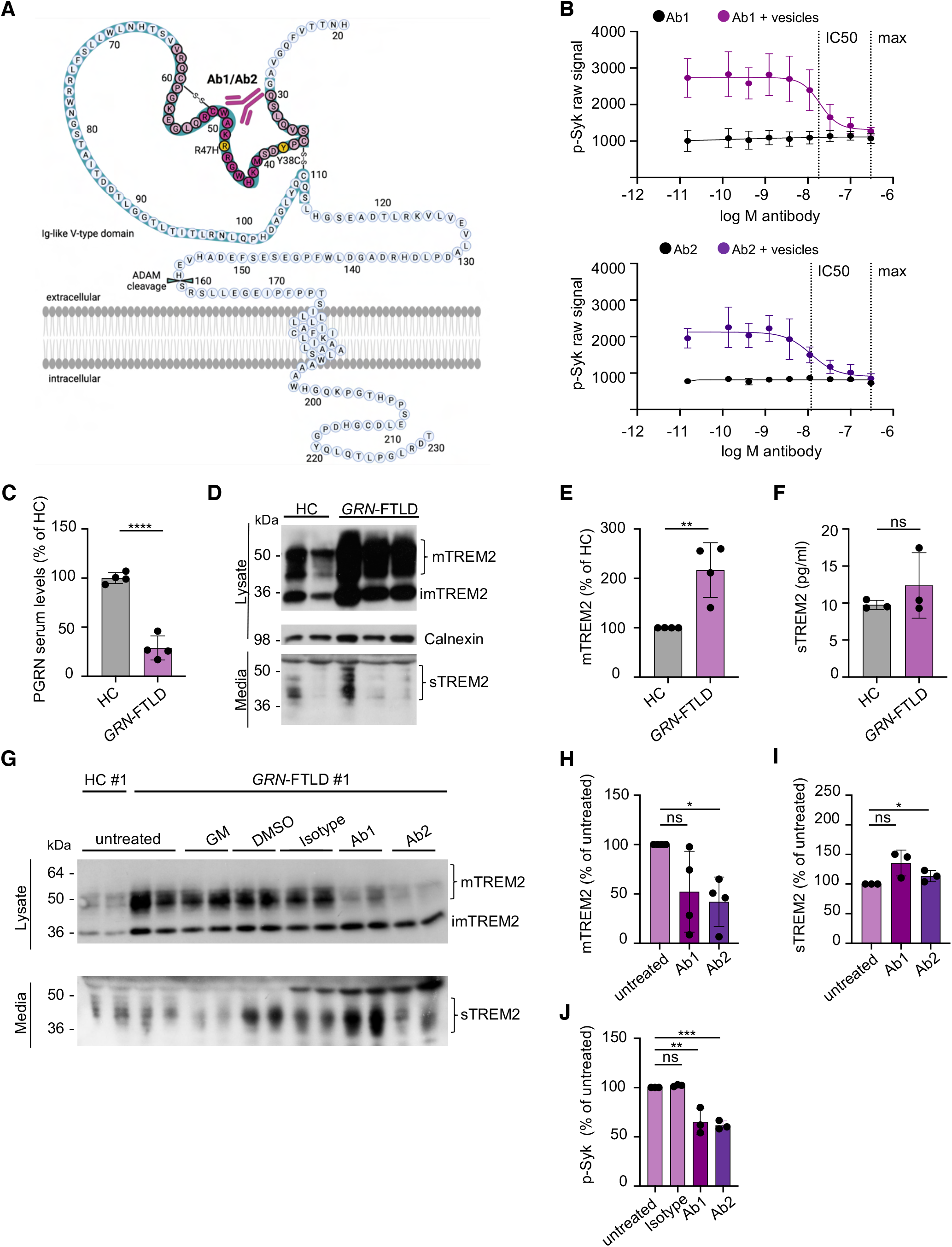
Human antagonistic TREM2 antibodies rescue elevated membrane-bound TREM2 levels and reduce p-Syk in primary human macrophages isolated from PGRN mutation carriers. **A.** Schematic presentation of human TREM2 with the identified binding site of antagonistic antibodies Ab1 and Ab2 (purple) in the Ig-like V-type domain. Light purple indicates the overlapping amino acid sequence of the two peptides, which are bound by Ab1 and Ab2 (see also EV1B). The disease associated Y38C and R47H mutations are marked in yellow. Created with BioRender.com **B.** AlphaLISA-mediated quantification of p-Syk in human macrophages with a dose titration treatment of Ab1 and Ab2 with or without liposomes for 5 min. IC50 and maximal inhibition (max) are indicated by a dashed line. Data represent the mean ± SEM (n=3 independent experiments). **C.** ELISA-mediated quantification confirms reduced PGRN serum levels in *GRN* mutation carriers vs. healthy controls. PGRN was measured by ELISA in technical triplicates and normalized to serum levels of healthy controls. Data points indicate individual patients (*GRN*-FTLD) and healthy controls (HC). **D.** Western blot of TREM2 in lysates and conditioned media of cultured human macrophages isolated from *GRN* mutation carriers and healthy controls. Mature (mTREM), immature (imTREM2), and soluble TREM2 (sTREM) are indicated. Calnexin was used as loading control. **E.** Quantification of mTREM2 expression levels in lysates of cultured human macrophages isolated from *GRN* mutation carriers (data shown in **D**). mTREM2 levels were normalized to healthy control (n=4). Data points indicate individual patients and healthy controls (HC). **F.** ELISA-mediated quantification of sTREM2 in conditioned media of human macrophages isolated from *GRN* mutation carriers and healthy controls (n=3). Isolated material from patient #3 did not yield enough material. Data points indicate individual patients and healthy controls (HC). **G.** Western blot of TREM2 in lysates and media of cultured human macrophages isolated from *GRN* mutation carrier #1 and healthy control #1 upon treatment with Ab1 and Ab2. An isotype antibody was used as a negative control. ADAM protease inhibition (GM) does not further increase mTREM2 levels in *GRN* mutation carriers. Equal amounts of protein were loaded. **H.** Quantification of mTREM2 expression normalized to healthy control (n=4) (data shown in **G**). Data points indicate individual patients and healthy controls (HC). **I.** ELISA-mediated quantification of sTREM2 in conditioned media of human macrophages isolated from PGRN mutation carriers and healthy controls (n=3). Isolated material from patient #3 did not yield enough material. Data points indicate individual patients and healthy controls (HC). **J.** AlphaLISA-mediated quantification of p-Syk levels in human macrophages upon treatment with Ab1 and Ab2 with liposomes for 60 min (n=3). An isotype antibody was used as a negative control. Data points indicate individual patients and healthy controls (HC). Isolated material from patient #3 did not yield enough material. Data represent mean ± SEM. For statistical analysis of samples the unpaired, two-tailed student’s t-test was performed. Statistical significance was set at *, *p* < 0.05; **, *p* < 0.01; ***, *p* < 0.001; ****, *p* < 0.0001; and ns, not significant

Next, we aimed to test if antagonistic TREM2 antibodies are capable of reducing TREM2 signaling in *GRN*-FTLD patient derived macrophages. To do so, we identified four patients with heterozygous *GRN* loss-of-function mutation (Fig EV2A). PGRN levels were determined in plasma of these patients and compared to healthy controls. This confirmed that all four *GRN* mutation carriers had significantly reduced plasma levels of PGRN (Fig 3C). We then generated monocyte-derived macrophages from peripheral blood samples of these patients and healthy volunteers. Western blot analysis revealed that macrophages of *GRN* mutation carriers show significantly enhanced levels of mature TREM2 as compared to healthy controls (Fig 3D and E). Although *GRN* mutation carriers express more mature TREM2 than healthy controls, sTREM2 in the conditioned media was not significantly altered (Fig 3D and F). Since evidence exists that shedding of TREM2 terminates cell autonomous signaling (Kleinberger et al., 2014; Schlepckow et al., 2017; Schlepckow et al., 2020; Thornton et al., 2017), these findings suggest that macrophages from *GRN-*FTLD patients exhibit increased TREM2 signaling, which occurs in conjunction with the hyperactivation microglial phenotype observed *in vitro* and *in vivo* (Gotzl et al., 2019).

Macrophages from *GRN*-FTLD patients and healthy controls were then treated with Ab1 and Ab2 for 24 h. Western blotting of cell lysates revealed that both TREM2 antibodies reduced mature TREM2, whereas an isotype control antibody had no effect (Fig 3G and H, Fig EV2B). The reduction of mature membrane bound TREM2 was accompanied by an increase of sTREM2 (Fig 3G and I). Thus, in line with the data shown in Fig 3A-B and Fig EV1A-G, both antibodies reduce signaling competent mature TREM2 and increase TREM2 shedding. To further demonstrate that TREM2 signaling can be modulated by TREM2 antagonistic antibodies in patient derived macrophages, we quantified Syk signaling. This demonstrated that both antagonistic antibodies reduce p-Syk in liposome stimulated macrophages, suggesting that antagonistic TREM2 antibodies may be capable to modulate TREM2 signaling in microglia in a beneficial manner (Fig 3J).

### Antagonistic TREM2 antibodies reduce hyperactivation of PGRN deficient human microglia

To corroborate and extend our findings in human myeloid cells we aimed to test modulation of TREM2 via our antagonistic antibodies in human induced pluripotent stem cells (iPSC) derived microglia (hiMGL). For this purpose, we generated *GRN* KO iPSC by targeting exon 2 using our established CRISPR genome editing pipeline ((Weisheit et al., 2020) see methods for details). We deeply phenotyped *GRN* KO iPSC to confirm loss of PGRN protein expression, maintenance of pluripotency, clonality as well as absence of unintended on- and off-target effects and chromosomal abnormalities (Weisheit et al., 2021) (Fig 4A-D; Fig EV3; Fig EV4). As expected, *GRN* KO hiMGL increased expression of cell surface TREM2 (Fig 4B and C) and showed enhanced shedding of TREM2 and consequently elevated levels of sTREM2 (Fig 4D). PGRN deficient hiMGL were treated with the antagonistic TREM2 antibodies described above. Consistent with the previously described results, antagonistic antibodies increased secretion of sTREM2 (Fig 4E). In line with this finding, both antagonistic antibodies reduced p-Syk signaling (Fig 4F). Moreover, both antibodies ameliorated the pathologically increased phagocytic activity of PGRN deficient hiMGL (Fig 4G), indicating that they dampen the activation state of PGRN deficient hiMGL. To further extend these findings, we asked if the antagonistic antibodies also correct the transcriptional signature of hyperactivated hiMGL. Therefore, we used a customized nCounter panel (NanoString Technologies) analyzing gene expression of 82 microglia-related genes and 8 housekeeping genes of WT and PGRN deficient hiMGL treated with the two antagonistic antibodies or isotype control (Fig 5A). Gene expression levels in each sample were normalized against the geometric mean of five housekeeping genes including *CLTC, HPRT1, RPL13A, TBP* and *PPIA*. DAM genes, such as *APOE*, *SPP1*, *CSF1*, *CCL3, LPL*, *TREM2, ITGAX* and *CD68*, were all significantly upregulated in PGRN deficient hiMGL compared to WT hiMGL (Fig 5A and B). Both antagonistic TREM2 antibodies significantly modulated the mRNA signature of PGRN deficient hiMGL towards a more homeostatic state (Fig 5A-D). Upregulation of TREM2 in PGRN deficient hiMGL was completely corrected by treating the cells with either antagonist antibody (Fig 5E). Upregulation of DAM genes was completely (*SPP1, CSF1, CCL3, LPL*, *ITGAX*) or at least partially (APOE and CCL2) rescued, while downregulation of the homeostatic marker *P2RY12* was reversed by antibody treatment (Fig 5F). Thus, TREM2 modulation with antagonistic antibodies ameliorates hyperactivation of microglia.

**Figure 4.**
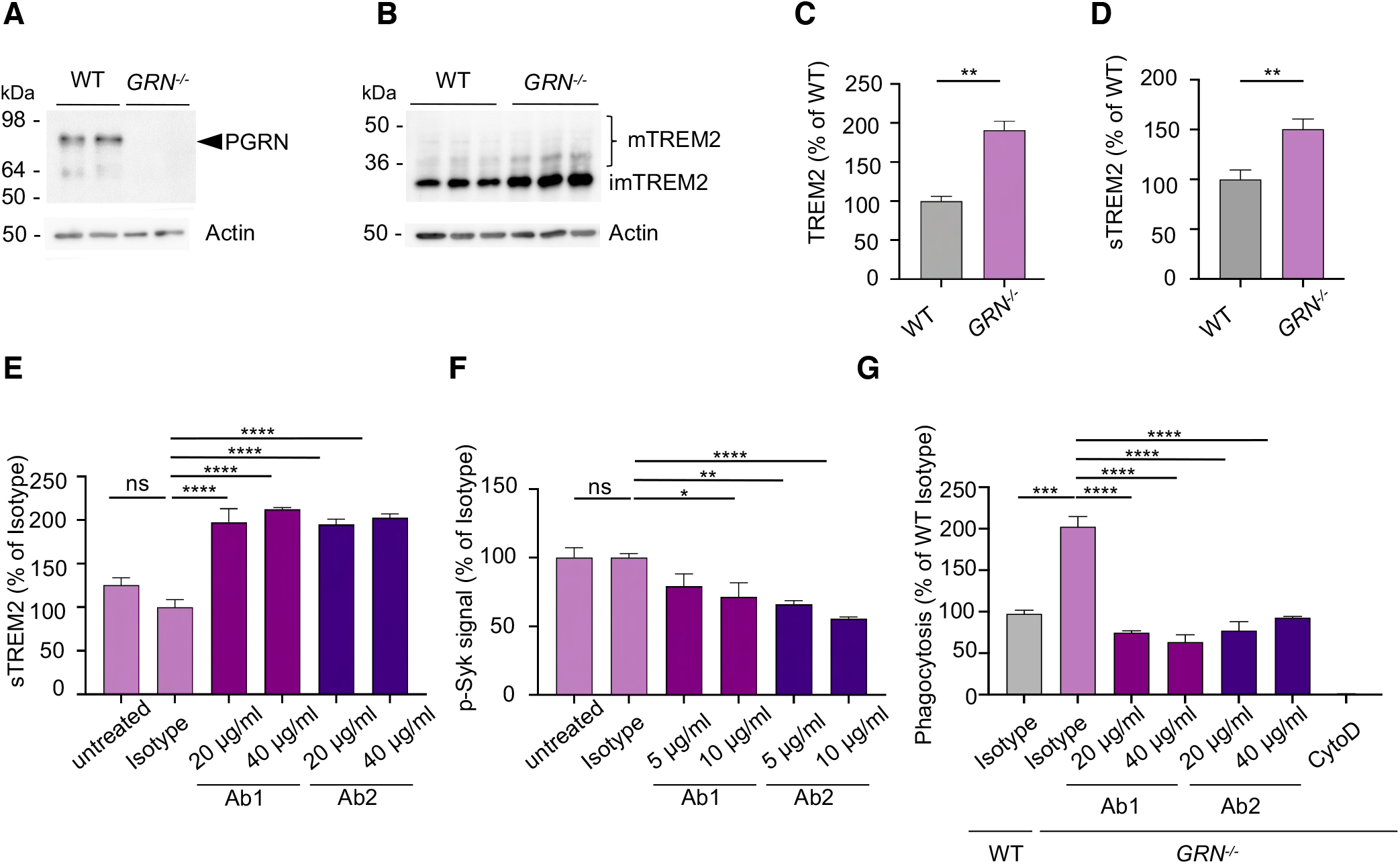
Antagonistic TREM2 antibodies reduce hyperactivation of PGRN deficient human microglia. **A.** Western blot of PGRN in whole-cell lysates of WT and *GRN^-/-^* hiMGL. Actin was used as loading control. **B.** Western blot of TREM2 in whole-cell lysates of WT and *GRN^-/-^* hiMGL. Mature (mTREM) and immature TREM2 (imTREM2) are indicated. Actin was used as loading control. **C.** Quantification of TREM2 expression in whole-cell lysates of WT and *GRN^-/-^* hiMGL (data shown in **B**). TREM2 levels were normalized to WT (n = 3). **D.** ELISA-mediated quantification of sTREM2 in conditioned media of WT and *GRN^-/-^* hiMGL (n = 3). **E.** ELISA-mediated quantification of sTREM2 in conditioned media of *GRN^-/-^* hiMGL upon treatment with Ab1 and Ab2 (20 μg/ml, 40 μg/ml) (n = 3). **F.** AlphaLISA-mediated quantification of p-Syk levels in *GRN^-/-^* hiMGL upon treatment with Ab1 and Ab2 (5 μg/ml, 10 μg/ml) with liposomes (1 mg/ml) for 5 min. An isotype antibody (10 μg/ml) was used as a negative control. (n = 8). **G.** Uptake assay for fluorescently labelled myelin. *GRN^-/-^* hiMGL phagocytose significantly more myelin as compare to WT hiMGL. This is reversed upon treatment with TREM2 antagonistic antibodies Ab1 and Ab2 (n=4). Data represent mean ± SEM. For statistical analysis in **C** and **D** the unpaired, two-tailed student’s t-test was performed, in **E** and **F** one-way ANOVA with Dunnett’s post hoc test, and in **G** one-way ANOVA with Tukey’s post hoc was used to compare untreated, Ab1, and Ab2 (20 µg/ml and 40µg/ml) conditions to the isotype treated condition. Statistical significance was set at *, *p* < 0.05; **, *p* < 0.01; ***, *p* < 0.001; ****, *p* < 0.0001; and ns, not significant.

**Figure 5.**
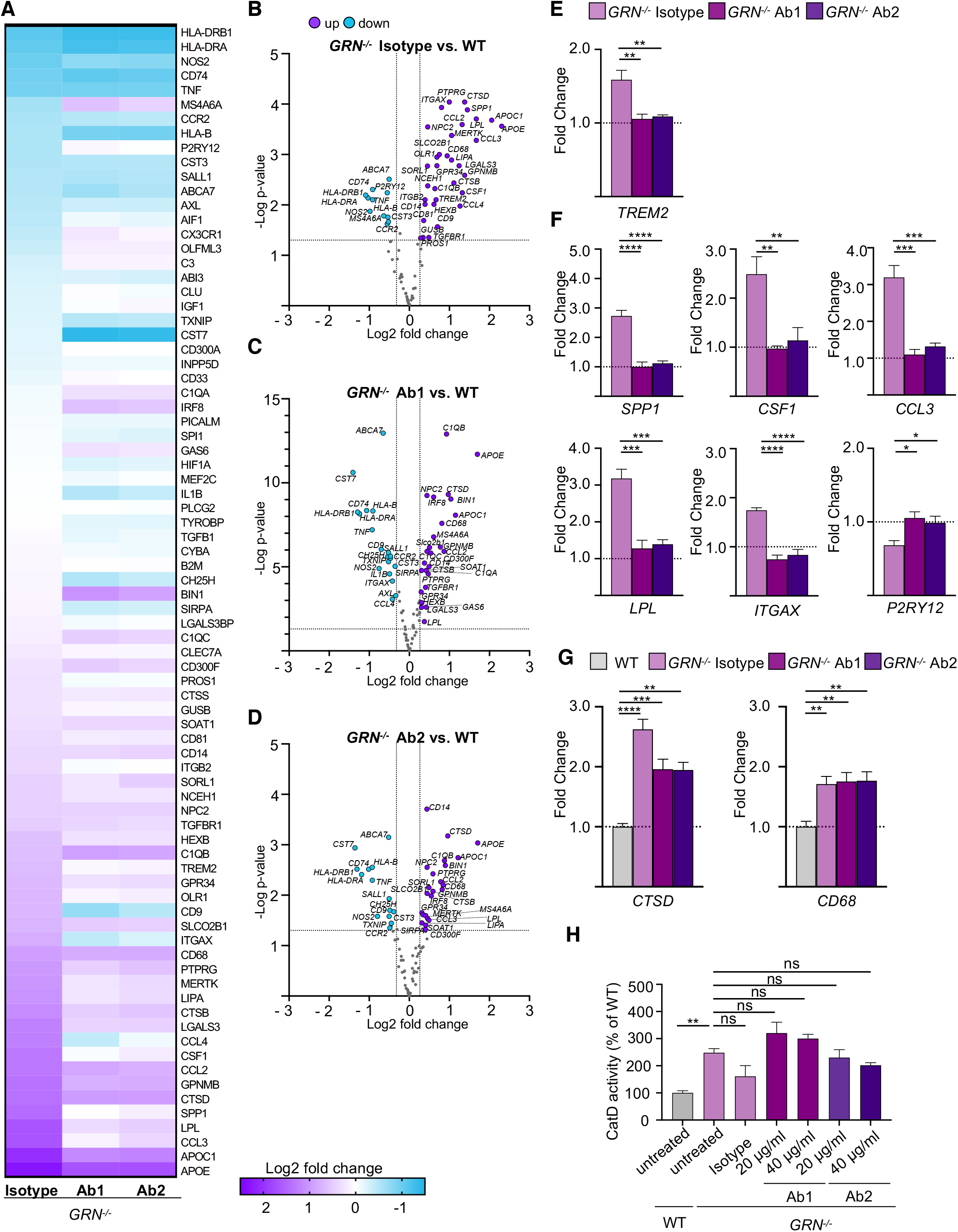
Antagonistic TREM2 antibodies reduce hyperactivation of PGRN deficient human microglia. **A.** Expression of all analyzed gene transcripts in *GRN^-/-^* hiMGL treated with control isotype antibody, Ab1 or Ab2 in comparison to WT hiMGL. Data show the mean of four individual NanoString measurements. The mRNA counts for each gene were normalized to the mean value of all WT samples followed by a log2 transformation. **B.** Volcano blot presentation of the differently expressed transcripts in *GRN^-/-^* hiMGL treated with isotype compared to WT hiMGL. Genes with more than 20% significantly changed expression are marked in purple (upregulated) or blue (downregulated). **C.** Volcano blot presentation of the differently expressed transcripts in *GRN^-/-^* hiMGL treated with Ab1 comparison to WT hiMGL. Genes with more than 20% significantly changed expression are marked in purple (upregulated) or blue (downregulated). **D.** Volcano blot presentation of the differently expressed transcripts in *GRN^-/-^* hiMGL treated with Ab2 comparison to WT hiMGL. Genes with more than 20% significantly changed expression are marked in purple (upregulated) or blue (downregulated). **E.** Transcript levels of *TREM2* in *GRN^-/-^* hiMGL treated with isotype control, antagonistic Ab1 and Ab2 normalized to the mean of the WT hiMGL samples. **F.** Levels of DAM gene transcripts significantly altered in *GRN^-/-^* hiMGL treated with Ab1 or Ab2 in comparison to isotype treatment from the data set in A normalized to the mean of the WT hiMGL samples. **G.** Transcript levels of *CTSD* and *CD68* of *GRN^-/-^* hiMGL treated with isotype control, Ab1 or Ab2 in comparison to WT hiMGL from the data set in A normalized to the mean of the WT hiMGL samples. **H.** Catalytic activity of cathepsin D (CatD) in untreated WT and *GRN^-/-^* hiMGL or *GRN^-/-^* hiMG treated with isotype control, Ab1 or Ab2 (20 μg/ml, 40 μg/ml), as measured by a CatD in vitro activity assay (n=3). Data represent mean ± SEM. For statistical analysis in **B** - **D** the unpaired, two-tailed student’s t-test was performed, in **E** and **F** one-way ANOVA with Dunnett’s post hoc test was used to compare Ab1 and Ab2 (20 µg/ml and 40µg/ml) conditions to the isotype treated condition, and in **G** - **H** one-way ANOVA with Dunnett’s post hoc test was used to compare Ab1, Ab2 (20 µg/ml and 40µg/ml) and isotype treated condition to WT cells. Statistical significance was set at *, *p* < 0.05; **, *p* < 0.01; ***, *p* < 0.001; ****, *p* < 0.0001, and ns, not significant.

### Reduced TREM2 signaling does not rescue lysosomal dysfunction

Next, we searched for a rescue of lysosomal phenotypes in PGRN deficient hiMGL. In contrast to the profound rescue of the homeostatic and diseases associated mRNA signatures upon treatment with the two antagonistic antibodies (Fig 5A-F), we did not observe a significant rescue of increased gene expression patterns associated with lysosomal dysfunction upon PGRN deficiency, like *CSTD*, *NPC2* and *CD68* mRNA expression (Fig 5A-D and G). Antagonistic antibodies also failed to rescue elevated cathepsin D (CatD) activity in PGRN deficient hiMGL (Fig 5H).

In total brain lysates of *Grn* KO and *Double* KO mice, CatD single chain (sc) and heavy chain (hc) were both increased (Fig 6A-C). Additionally, in brain lysates from 14-months-old *Double* KO mice no reduction of enhanced CatD expression could be observed (Fig 6A-C). Furthermore, the catalytic activity of CatD, which was increased in *Grn* KO mice in an age dependent manner (Fig 6D and E) was also not rescued by the additional loss of TREM2 (Fig 6E), suggesting that lysosomal dysfunction of *Grn* KO mice cannot be rescued by TREM2 modulation. To further support this, we investigated *Grn* KO brains for the accumulation of lipofuscin, an autofluorescent lipopigment found in several lysosomal storage disorders (Gotzl et al., 2014). In line with the failure of the *Double* KO to rescue lysosomal hyperactivity, lipofuscin accumulation was not reduced upon loss of TREM2 in *Grn* KO mice although surprisingly almost no lipofuscin was observed in single *Trem2* KO mice (Fig 6F and G).

**Figure 6.**
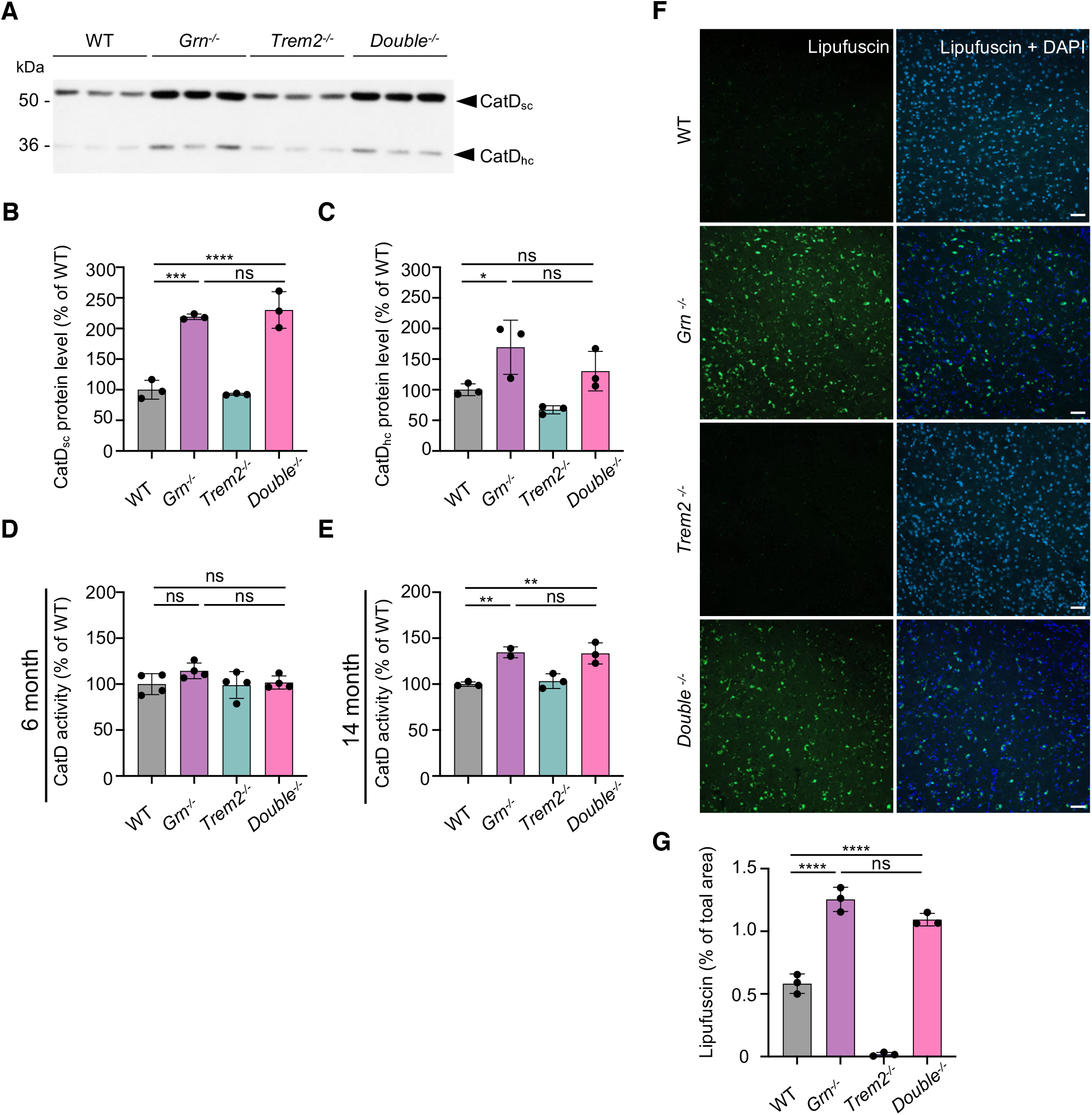
Abolishing TREM2 signaling does not rescue lysosomal dysfunction in *Grn*^-/-^ mice. **A.** Western blot of CatD in total brain lysates from 14-month-old WT, *Grn^-/-^, Trem2^-/-^, and Double^-/-^* mice. CatD maturation variants are indicated (sc: single chain; hc: heavy chain; n=3). **B., C.** Quantification of CatD variants in **A** normalized to WT. **D., E.** Catalytic activity of CatD in brain lysates from 6-month-old (**D**) or 14-month-old (**E**) *Grn^-/-^, Trem2^-/-^, Double^-/-^* mice normalized to WT (n=3 per genotype). **F.** Immunohistochemical analysis of lipofuscin (green) in coronal brain sections. Representative images of hypothalamus. Scalebars = 50 μm. **G.** Quantification of lipofuscin autofluorescence. Five pictures per mouse were taken, means were normalized to WT samples (n=3 per genotype). Data represent mean ± SEM. For statistical analysis one-way ANOVA with Tukey’s post hoc test of *Grn^-/-^, Trem2^-/-^* and *Double^-/-^* was used. Statistical significance was set at *, *p* < 0.05; **, *p* < 0.01; ***, *p* < 0.001; ****, *p* < 0.0001, and ns, not significant.

### Loss of TREM2 does not rescue lysosomal lipid dyshomeostasis in *Grn* KO mice

Previous studies have examined the impact of either *Trem2* or *Grn* deletion on the lipidome of mouse brain. In the case of *Trem2*, no significant lipid changes were observed in the *Trem2* KO mouse brain at baseline, although upon cuprizone challenge a striking accumulation of cholesterol esters and various sphingolipids was revealed (Nugent et al., 2020). In the *Grn* KO mouse, a recent study described an age-independent deficit in levels of the lysosomal lipid bis(monoacylglycerol)phosphate (BMP) that was accompanied by an age-dependent accumulation of the GCase substrate glucosylsphingosine (GlcSph) (Logan et al., 2021). To determine whether deletion of *Trem2* on the *Grn* KO background has any effect on the composition of the brain lipidome, we performed targeted lipidomic analysis using LCMS on 6-month-old WT, *Grn* KO, *Trem2* KO and *Double* KO mouse brain homogenates (Fig 7). As previously described (Nugent et al., 2020), the *Trem2* KO showed no significant differences in brain lipid content relative to WT mice (Fig 7B), while the *Grn* KO showed a significant decrease in several BMP species as well as an increase in GlcSph (Fig 7A and E-G), which is consistent with previous data (Logan et al., 2021). Consistent with previous findings (Arrant et al., 2019; Logan et al., 2021; Zhou et al., 2019) and the increased accumulation of the GCase substrate GlcSph, we found a significant decrease in the GCase activity in *Grn* KO mice and *Double* KO (Fig 7H). Importantly, genetic interaction analysis demonstrated no statistically significant difference in the levels of any analyte in the *Double* KO brain compared to the *Grn* KO alone (Fig 7D). Thus, reduction of TREM2 fails to correct lysosomal dysfunction and abnormal lipid metabolism in PGRN deficient mice.

**Figure 7.**
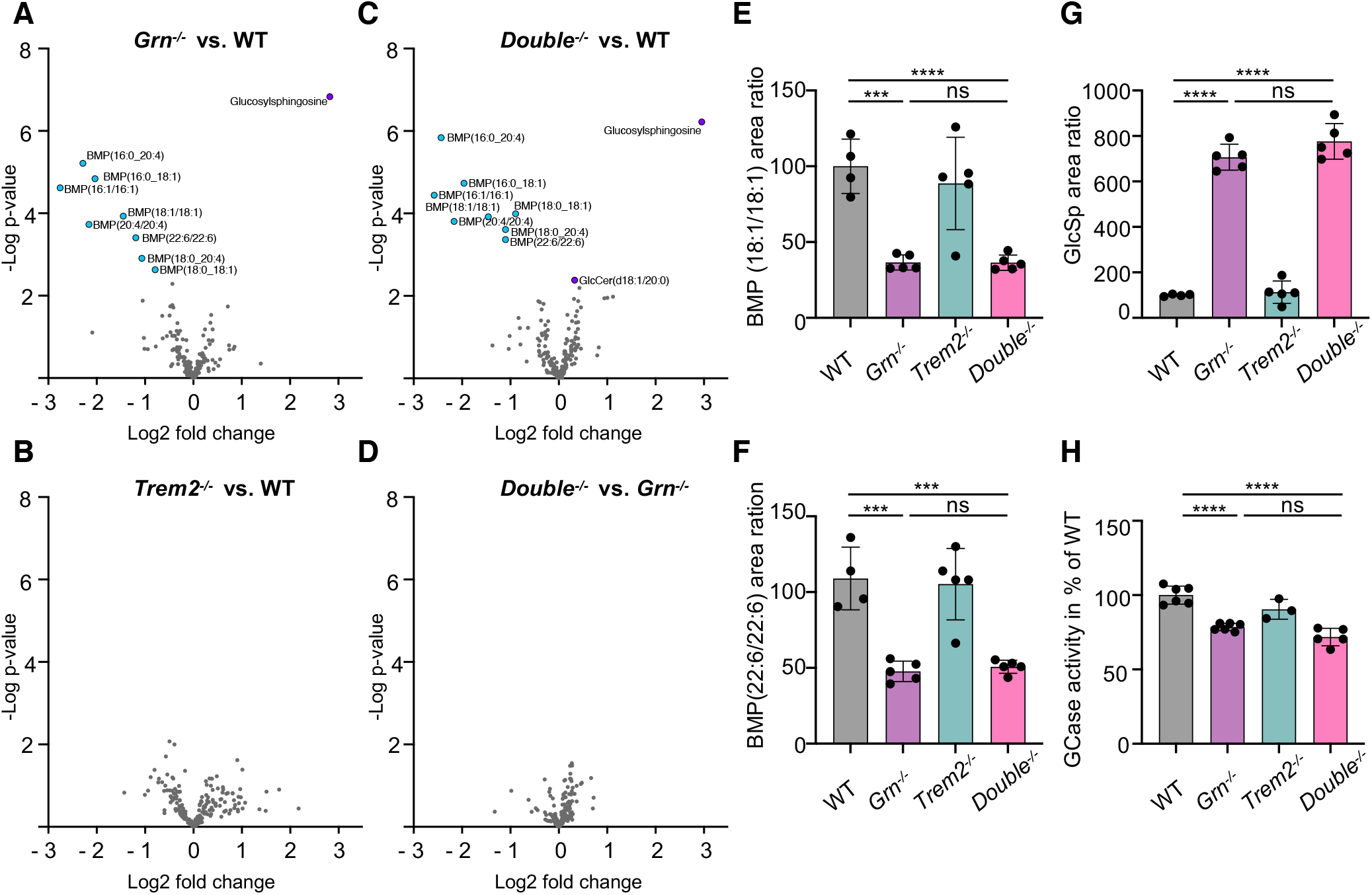
Reduced TREM2 signaling does not rescue dysregulated lipids in *Grn^-/-^* mice. **A-D.** Volcano plot presentation of lipids and metabolites upregulated (purple) or downregulated (blue) in total brain homogenates from 6-month-old *Grn^-/-^* (A)*, Trem2^-/-^* (B)*, Double^-/-^* (C) mice in comparison to WT, and *Double^-/-^* in comparison to *Grn^-/-^* mice (D). Counts for each sample were normalized to the mean value of WT followed by a log2 transformation (n=4-5 per genotype). Analyte values were adjusted with an FDR <10% to exclude type I errors in null hypothesis testing. **E-G.** Abundance of BMP species and glucosylsphingosine (GlcSph) in total brain of 6-month-old *Grn^-/-^, Trem2^-/-^, Double^-/-^* and WT mice (n=4-5 per genotype). **H.** Glucocerebrosidase (GCase) activity in whole brain lysates from 6-month-old *Grn^-/-^, Trem2^-/-^, Double^-/-^* and WT mice. The linear increase of fluorescence signal was measured and then normalized to WT mice (n=3-6 per genotype). Data represent mean ± SEM. For statistical analysis one-way ANOVA with Tukey’s post hoc test of *Grn^-/-^, Trem2^-/-^* and *Double^-/-^* was used. Statistical significance was set at *, *p* < 0.05; **, *p* < 0.01; ***, *p* < 0.001; ****, *p* < 0.0001, and ns, not significant.

### Enhanced brain pathology in double knockout mice indicates a protective function of hyperactivated microglia

The above-described findings suggest that lowering TREM2 reduces hyperactivation of microglia. Although based on our extensive gene expression analyses there is no indication of an enhanced inflammatory response or upregulated complement factor expression, in the *Double* KO no rescue of lysosomal deficits was observed. To prove if reduction of microglial hyperactivation ameliorates secondary neurodegeneration, we analyzed the concentrations of neurofilament light chain (NfL), a fluid biomarker for neuronal damage (Preische et al., 2019), in the cerebrospinal fluid (CSF) of 14-month-old mice. In line with previous findings (Zhang et al., 2020), NfL was increased in PGRN deficient mice whereas no change was observed in *Trem2* KO animals as compared to WT mice (Fig 8A). Surprisingly, we found a striking increase of NfL in the *Double* KO mice (Fig 8A), suggesting that reduced microglial hyperactivation in PGRN deficient mice increases progression of neurodegeneration. To further elucidate which genes and pathological pathways may be ameliorated or enhanced by lowering TREM2 in PGRN deficient mice, we isolated mRNA from total brain lysates of all three mouse models and searched for changes in mRNA expression using the nCounter Neuropathology panel (NanoString Technologies) (Werner et al., 2020). The neuropathology panel with 770 genes included was specifically designed to analyze neurodegenerative phenotypes in mouse models and allows investigating six fundamental themes of neurodegeneration, namely neurotransmission, neuron-glia interaction, neuroplasticity, cell structure integrity, neuroinflammation and metabolism. Analysis of total brain mRNA confirmed the rescue of the *Grn* KO associated DAM signature in the *Double* KO mice (Fig EV5A-E) and revealed compared to WT mice no significant upregulation to genes associated to neuroinflammation like *Gfap*, *Tnf*, or *Tnfrsf11b* or to genes associated to synaptic pruning, such as the complement factors (*C1qc*, *C1qa*, *C1qb*) (Fig EV5E and F). Strikingly, compared to single *Grn* KO and *Trem2* KO only two genes were significantly altered and downregulated in the *Double* KO: the transcription factor *Npas4* (Neuronal PAS domain protein 4), which regulates activation of genes involved in the excitatory-inhibitory balance and is known to exert neuroprotective activities (Fu et al., 2020; Spiegel et al., 2014) (Fig 8C; Fig EV5E and F), and *Grn3b* (Perez-Otano et al., 2016), a glutamate receptor subunit (Fig 8C; Fig EV5E and F). Pathway analysis in *Grn* KO mice revealed the highest increases in “autophagy”, “activated microglia”, “angiogenesis” and “disease association” associated clusters (Fig 8B). These four pathways score very low in *Trem2* KO mice, again confirming opposite effects of the two single KOs. All four pathways show a lower score in the *Double* KO than in the single *Grn* KO and three of these pathways namely, “activated microglia”, “angiogenesis” and “disease association” are downregulated compared to WT. However, other pathways like “neuronal cytoskeleton”, “tissue integration”, and “transmitter synthesis and storage, transmitter response and uptake”, are most heavily affected in the *Double* KO, which is consistent with enhanced neuropathological phenotypes.

**Figure 8.**
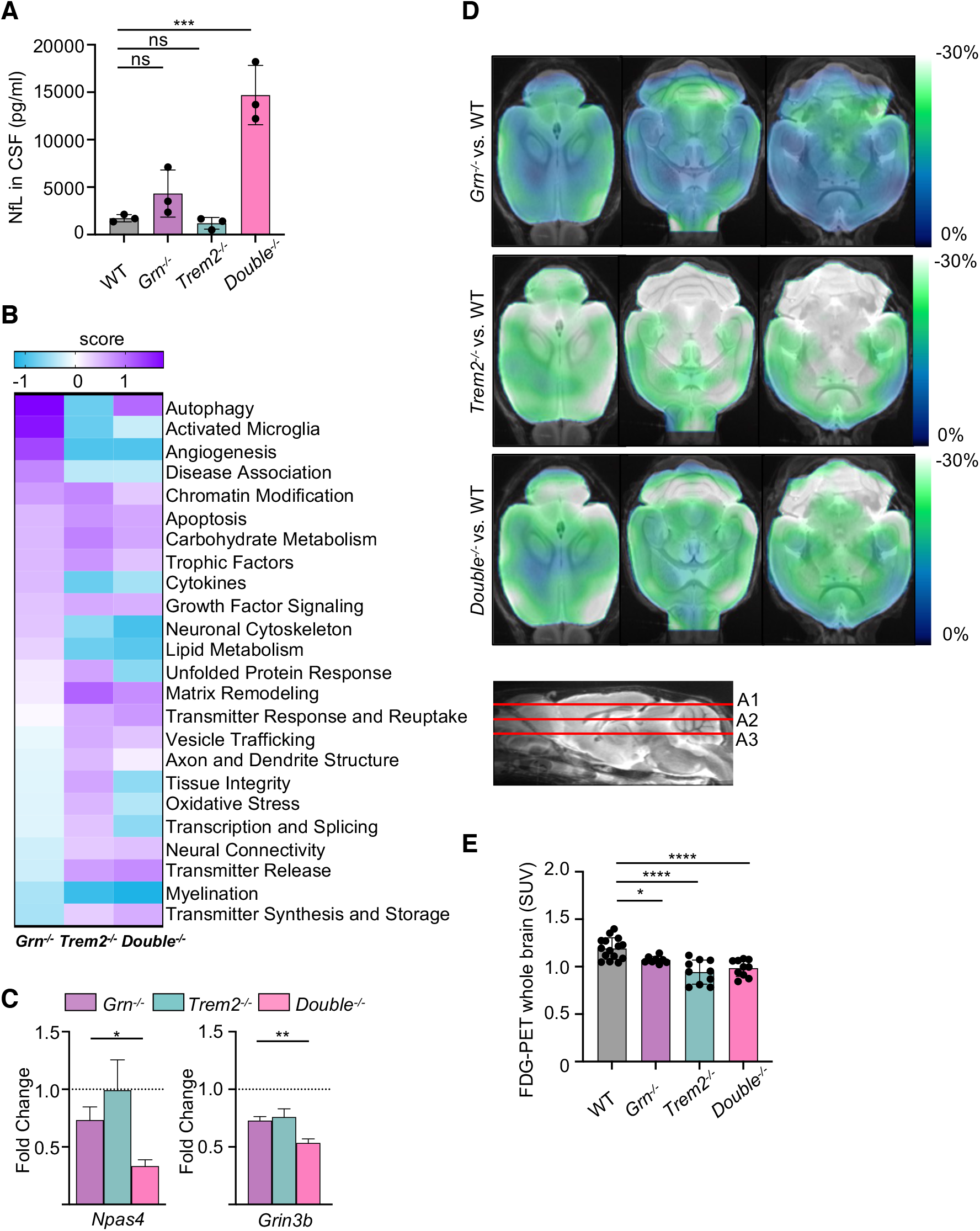
Hyperactivation of microglia in *Grn*^-/-^ mice is not deleterious. **A.** Immunoassay-based quantification of neurofilament-light (NfL) levels in CSF of 14-month-old *Grn^-/-^, Trem2^-/-^, Double^-/-^* and WT mice (n=3). **B.** Neuropathology NanoString panel analysis of total brain mRNA expression of 6-month-old *Grn^-/-^, Trem2^-/-^, Double^-/-^* and WT mice based on NanoString advanced analysis R-script included in the panel (n=4). **C.** Expression levels of *Npas4* and *Grin3b* of *Grn^-/-^, Trem2^-/-^* and *Double^-/-^* microglia from the data set in A. mRNA expression levels are normalized to the mean of the WT cohort. **D.** Axial slices as indicated show %-FDG-PET differences between *Grn^-/-^, Trem2^-/-^* and *Double^-/-^* (all cold colour scales) when compared to WT at the group level. Images were adjusted to an MRI template. **E.** Bar graph illustrates individual FDG-PET values derived from a whole brain volume of interest. Data represent mean ± SD. 8-15 female mice per group at an average age of 10.7 ± 1.5 months (*Grn^-/-^* n = 8, *Trem2^-/-^* n = 10, *Double^-/-^* n = 10, WT n = 15) were used. Data represent mean ± SEM. For statistical analysis in **A** one-way ANOVA with Dunnett’s post hoc test was used the unpaired, in **C** two-tailed student’s t-test was performed, and in **E,** one-way ANOVA with Tukey’s post hoc test was used. Statistical significance was set at *, *p* < 0.05; **, *p* < 0.01; ***, *p* < 0.001; ****, *p* < 0.0001, and ns, not significant.

Since the above-described findings suggest that brain function of PGRN deficient mice may not be improved by the additional knockout of TREM2, we investigated if the additional loss of TREM2 in PGRN deficient mice ameliorates deficits in glucose uptake *in vivo*. To determine brain cerebral uptake rates of glucose in *Double* KO in vivo, we performed 2-[18F]fluoro-d-glucose PET (FDG-PET). We confirmed a reduced cerebral glucose uptake in *Grn* KO (p < 0.05) and *Trem2* KO (p < 0.0001) mice compared to WT (Fig 8D and E) as described in previous studies (Gotzl et al., 2019; Kleinberger et al., 2017). However, we still observed similar decreased glucose uptake in *Double* KO mice when compared to WT (p < 0.0001) (Fig. 8D and E), revealing a consistently decreased glucose uptake between *Grn* KO, *Trem2* KO or *Double* KO mice and WT mice.

Together, these findings indicate that reducing hyperactivation of microglia does not ameliorate lysosomal dysfunction of PGRN deficient mice but may even promote neurodegeneration.

## Discussion

PGRN and the proteolytically derived granulin peptides may have important lysosomal functions, as exemplified by the identification of homozygous loss-of-function *GRN* mutations, which are causative for NCL (Smith et al., 2012). Accumulating evidence suggests that PGRN /granulins directly or indirectly regulate the activity of lysosomal enzymes such as CatD, (Beel et al., 2017; Butler et al., 2019a; Butler et al., 2019b; Valdez et al., 2017; Zhou et al., 2017), GCase (Arrant et al., 2019; Jian et al., 2016; Zhou et al., 2019), and HexA (Chen et al., 2018). The last two enzymes are involved in sphingolipid degradation, a process regulated by the lysosomal phospholipid BMP, which is stabilized by PGRN (Logan et al., 2021; Paushter et al., 2018; Schulze and Sandhoff, 2014). PGRN may affect lysosome acidification and thereby lysosomal enzyme activity (Logan et al., 2021; Tanaka et al., 2017). We and others have shown that PGRN deficiency results in upregulation of several lysosomal enzymes (Gotzl et al., 2018; Gotzl et al., 2014; Klein et al., 2017; Root et al., 2021). However, it remained unclear if microglial hyperactivation observed in PGRN deficient microglia contributes or is a consequence of lysosomal dysfunction. Activated microglia are found in late stages of many neurodegenerative diseases including AD and FTLD, and are known to be deleterious by promoting synaptic pruning and neuronal cell death (Heneka et al., 2013; Hong et al., 2016a; Hong et al., 2016b). Specifically, FTLD patients suffering from *GRN* haploinsufficiency show pathological hyperactivation of microglia as measured by TSPO-PET (Gotzl et al., 2019; Marschallinger et al., 2020; Martens et al., 2012; Zhang et al., 2020). Similarly, mice lacking PGRN exhibit hyperactivation of microglia as indicated by an enhanced DAM signature including TREM2, an increased TSPO signal, and increased phagocytic and synaptic pruning activity (Gotzl et al., 2019; Lui et al., 2016; Zhang et al., 2020). We therefore asked if the pathological outcome of PGRN deficiency may be promoted by TREM2-dependent microglial overaction. To address this question, we sought to reduce TREM2 dependent signaling by two independent strategies: genetic loss-of-function and pharmacological inhibition with antagonist antibodies. To achieve the former, we crossed *Trem2* KO to the *Grn* KO to generate a *Double* KO mouse. For the latter approach, we identified TREM2 antagonistic antibodies, which negatively regulate TREM2 by increasing surface receptor shedding and preventing lipid ligand induced signaling of the co-receptor DAP12. Both approaches successfully dampened several aspects of TREM2 dependent microglial activation. However, although reduction of TREM2 signaling by two independent approaches lessened microglial hyperactivation to some extent, this was not sufficient to ameliorate lysosomal deficits, dysregulation of lysosomal lipids, and reduced glucose uptake. These findings demonstrate that microglial hyperactivation is secondary to the primary loss of lysosomal function caused by PGRN deficiency. Surprisingly, inhibition of TREM2 function results in changes in markers for neurodegeneration and synapse loss in *Double* KO animals. Our extensive gene expression analyses do not imply that the total loss of TREM2 function in *Grn* KO mice causes additional neurotoxicity for example by supporting pro-inflammatory microglial populations. Thus, the fact that the additional loss of TREM2 leads to increased brain pathology, indicates that TREM2-regulated microglial activation states may not necessarily be deleterious and may in fact be protective. We suggest that hyperactivated microglia e.g., in *Grn* KO mice, resemble the previously described DAM2 microglia or may develop into them by even further increasing their DAM signature (Keren-Shaul et al., 2017). Consistently, fully activated DAM2 microglia were recently described to be particularly protective in a mouse model for amyloidosis and tau pathology (Lee et al., 2021). This is very surprising since chronically activated microglia, as observed in PGRN loss-of-function models and mouse models for amyloid and tau pathology would have been expected to exert significant damage within the brain, for example by induction of the inflammasome (Heneka et al., 2018). However, our findings together with those by Lee et al. (Lee et al., 2021) rather suggest that TREM2 dependent chronic activation is protective, which may have implications for therapeutic attempts employing modulation of TREM2 activity by agonistic antibodies (Deczkowska et al., 2020; Lewcock et al., 2020). In that regard the nomenclature used for describing diverse microglial states, namely homeostatic, disease associated, and hyperactivated, may also be reconsidered. We find these terms misleading, since they indicate that homeostatic microglia are beneficial whereas disease associated or hyperactivated microglia are deleterious. Importantly, the brain environment and pathological context is important for ascribing microglial state and associated functions which requires deeper understanding beyond transcriptional characterization to elucidate the impact to brain function. However, one may describe these fundamentally different populations of microglia as “surveilling” versus “responding” microglia. The term “responding” would implicate that these microglia exert protective effects. Protective microglial functions are promoted by enhancing TREM2 signaling with agonist TREM2 antibodies (Lewcock et al., 2020). All currently described agonistic TREM2 antibodies act via similar mechanisms by inhibiting shedding and directly activating TREM2, and therefore increasing functional receptor on the cell surface (Lewcock et al., 2020). Notably, all known agonistic TREM2 antibodies bind to the stalk region close to the cleavage site by ADAM10/17 (Lewcock et al., 2020; Schlepckow et al., 2017). In contrast the two antagonistic antibodies described here bind in the IgV-fold between amino acids 30 and 63 (Fig A). Interestingly, this domain of TREM2 harbors a number of AD and FTLD associated sequence variants (Colonna and Wang, 2016). The R47H variant increases AD risk and affects TREM2 dependent microglial proliferation, lipid metabolism and microgliosis. Similarly, the FTLD associated Y38C variant causes a loss-of-function by misfolding and retention of TREM2 within the endoplasmic reticulum (Kleinberger et al., 2014). Thus, the antagonistic antibodies appear to bind at a functionally critical region and may displace natural ligands (*e.g.*, lipids), thus preventing induction of TREM2 signaling, in addition to promoting TREM2 shedding.

Taken together, eliminating TREM2 function by two independent approaches, and thus reducing hyperactivation, does not rescue lysosomal dysfunction caused by GRN deficiency, but rather exacerbates pathological endpoints characteristic for neurodegeneration, including elevation of CSF NfL and reduced transcription of the neuroprotective transcription factor *Npas4*. Thus, hyperactivated microglia retain against common assumptions TREM2 dependent protective functions.

## Materials and Methods

### Animal experiments and mouse brain tissue

All animal experiments were performed in accordance with German animal welfare law and approved by the government of upper Bavaria. Mice were kept under standard housing conditions including standard pellet food and water provided *ad libitum*. Mice were sacrificed by CO_2_ inhalation or deep/lethal anesthesia followed by PBS perfusion. Brain tissue was obtained from male and female of the following mouse strains: C57BL/6J *Grn* (Kayasuga et al., 2007) and *Trem2* knockout line (Turnbull et al., 2006). PET scans and CSF withdrawal were performed under the animal license: ROB 55.2-2532.Vet_02-18-32.

### Isolation, differentiation and culture of human primary monocytes

Human primary monocytes were isolated from whole blood using Sepmate tubes (StemCell Technologies, #85450) in combination with RosetteSep Human Monocyte Enrichment Cocktail (StemCell Technologies, #15068) according to the manufacturer’s protocol. Briefly, fresh blood was collected into EDTA-coated collection tubes and stored at room temperature (RT) until further processing for maximal 6 h. EDTA was added to a final concentration of 1 mM and tubes were mixed by inversion. 50 μl/ml blood of RosetteSep cocktail was added and samples for incubated for 20 min at RT. Cells were separated on a density gradient using Ficoll-Paque PLUS (ThermoFisher, #11768538). After centrifugation isolated cells were washed with PBS supplemented with 2% FCS. Leftover red blood cells were lysed using ACK lysing buffer (ThermoFisher, #11509876) for 2 min at RT. Subsequently, cells were washed two times with PBS supplemented with 2% FCS. Cells were counted using Trypan blue as a viability dye and 1 x 10^6^ cells were plated in 10 cm dishes in 10 ml RPMI medium supplemented with 10 % FCS, 10 % NEAA, 10 % L-glutamine, 10 % sodium pyruvate and M-CSF with a final concentration of 50 ng/ml. 48 h after isolation 1 ml medium with 500 ng/ml M-CSF was added to the cells. Five days after isolation, cells were washed with PBS and scraped. Cells were counted as described above and plated in either 96-well plates at a density of 5×10^4^ cells/well in 100 μl or in 12-well plates at a density of 3×10^5^ cells/well in 600 μl RPMI medium supplemented with 10 % FCS, 10 % NEAA, 10 % L-glutamine, 10 % sodium pyruvate and M-CSF with a final concentration of 50 ng/ml.

### Generation and maintenance of *GRN* KO iPSC lines

iPSC experiments were performed in accordance with all relevant guidelines and regulations. Female iPSC line A18944 was purchased from ThermoFisher (#A18945). iPSCs were grown in Essential 8 Flex Medium (ThermoFisher, #A2858501) on VTN-coated (ThermoFisher, #A14700) cell culture plates at 37°C with 5% CO_2_ and split twice a week as small clumps after a 5 min incubation in PBS/EDTA. Prior to electroporation, iPSCs were split to single cells after a 10 min incubation in PBS/EDTA to Geltrex-coated (ThermoFisher, #A1413302) plates and cultured in StemFlex Medium (ThermoFisher, #A3349401) containing 10 mM ROCK inhibitor (Selleck-chem S1049) for two days. iPSCs were transfected by electroporation as described earlier (Kwart et al., 2017) with modifications. Briefly, two million cells were harvested with Accutase (ThermoFisher, #A1110501), resuspended in 100 ml cold BTXpress electroporation solution (VWR International GmbH, #732-1285) with 20 mg Cas9 (pSpCas9(BB)-2A-Puro (PX459) V2.0 (gift from Feng Zhang; Addgene plasmid #62988; http://n2t.net/addgene:62988; RRID: Addgene_62988 (Ran et al., 2013)) and 5 mg sgRNA cloned into the BsmBI restriction site of the MLM3636 plasmid (gift from Keith Joung, Addgene plasmid #43860; http://n2t.net/addgene:48360; RRID: Addgene_43860). Cells were electroporated with 2 pulses at 65 mV for 20 ms in a 1 mm cuvette (ThermoFisher, #15437270). After electroporation, cells were transferred to Geltrex-coated 10 cm plates and grown in StemFlex Medium containing 10 mM ROCK inhibitor until visible colonies appeared. Cells expressing Cas9 were selected by 350 ng/ml Puromycin dihydro-chloride (VWR International GmbH, #J593) for three consecutive days starting one day after electroporation (Steyer et al., 2018). Single-cell clone colonies were then picked and analyzed by RFLP assay, using NEB enzyme MwoI for the *GRN* KO, and Sanger sequencing as previously described (Kwart et al., 2017).

#### CRISPR/Cas9 genome editing

Design and preparation of editing reagents and quality control of edited iPSCs was performed as described previously (Weisheit et al., 2020; Weisheit et al., 2021). Briefly, we used CRISPOR (http://crispor.tefor.net, (Concordet and Haeussler, 2018) to select guide RNAs and determine putative off-target loci. We chose gRNAs targeting exon 2 of *GRN*, as it is present in most splice isoforms and a frameshift would affect large parts of the coding region. We also ensured presence of nearby stop codons in alternate reading frames in the sequence after the cut site. Successful knockout was confirmed on mRNA level by qPCR, and on protein level by Western blot using RIPA lysate and ELISA using conditioned media, respectively. For quality control of edited iPSC clones, we checked absence of off-target effects by PCR-amplifying and Sanger sequencing the top 5 hits based on MIT and CFD scores on CRISPOR. We also confirmed absence of on-target effects such as large deletions and loss of heterozygosity using qgPCR and nearby SNP sequencing (Weisheit et al., 2021). Finally, we also ensured pluripotency by immunofluorescence staining for typical markers OCT4, NANOG, SSEA4 and TRA160, and chromosomal integrity by molecular karyotyping (LIFE & BRAIN GmbH).

#### Differentiation of human iPSC-derived Microglia (hiMGL)

We differentiated hiMGL from iPSCs as described (Abud et al., 2017) with modifications to improve efficiency and yield: When iPSCs were 70-90% confluent, they were split 1:100-200 onto GelTrex-coated 6-well plates for the HPC differentiation using EDTA to get around ∼20 small colonies per well. Cells were fed with 2 ml of HemA medium (HPC differentiation kit, StemCell Technologies) on day 0 and half-fed with 1 ml on day 2. Media were switched to 2 ml of HemB on day 3 with half-feeds on days 5 and 7 and 1 ml added on top on day 10. On day 12 HPCs were collected as non-adherent cells to either freeze or continue with the microglia differentiation. HPCs were frozen at 1 million cells per ml in BamBanker (FUJIFILM Wako Chemicals). They were then thawed directly onto GelTrex-coated 6-well plates with 1 million cells evenly distributed among 6 wells in 2 ml iMGL media with 25 ng/ml M-CSF, 100 ng/ml IL-34, and 50 ng/ml TGF-ß added fresh. 1 ml of media was added every other day. During the microglia differentiation, the cells were split 1:2 every 6-8 days depending on confluency. A very similar differentiation protocol was published recently (McQuade et al., 2018). We did do not use CD200 and CX3CL1 as this did not have an effect on hiMGL gene expression, as determined by NanoString analysis (data not shown). hiMGL were used for experiments on day 16 of the differentiation.

### Antagonist antibody generation and verification

Antibody generation was carried out by performing single B-cell sequencing on lymphoid tissues from rodents immunized full length human TREM2 ectodomaine (ECD)-Fc protein (AbCellera Inc). Antibodies were screened based on binding to human TREM2, and clones of interest were reformatted onto human effectorless human IgG1-LALAPG backbones for material generation and further evaluation cell binding potency and functional impact to TREM2 signaling. Antagonists were identified by their ability to block lipid ligand-induced activation of p-Syk on HEK293 cells over-expressing TREM2-DAP12.

#### Affinity determination and binding kinetics

Human TREM2 binding affinities of anti-TREM2 antibodies were determined by surface plasmon resonance using a Biacore 8K instrument. Biacore Series S CM5 sensor chip was immobilized with a mixture of two monoclonal mouse anti-Fab antibodies (Human Fab capture kit from GE Healthcare) to capture antibodies for binding measurements. In order to measure human TREM2 binding affinities of anti-TREM2 antibodies, serial 3-fold dilutions of recombinant human TREM2-ECD protein were injected at a flow rate of 30 μl/min for 300 seconds followed by 600 second dissociation in HBS-EP+ running buffer (GE, #BR100669). A 1:1 Languir model of simultaneous fitting of k_on_ and k_off_ was used for antigen binding kinetics analysis.

#### Epitope mapping of antagonist TREM2 antibodies

Biotinylated polypeptides for human Trem2 IgV domain (Sequences in Fig EV1B) were purchased from Elim Biopharmaceuticals, Inc.. N-terminal cysteine was added to peptides to enable maleimide-thiol conjugation of biotin. The lyophilized biotinylated peptides were reconstituted in 20 mM Tris buffer at pH 8.0. Antibody binding to TREM2 IgV domain peptides was detected using a sandwich ELISA. Briefly, a 96 well half-area ELISA plate was coated with streptavidin overnight at 4°C. The following day, biotinylated TREM2 IgV peptides diluted to 1 µM in 1% BSA/PBS were added to the plate and incubated for 1 h. Antibodies diluted to 120 nM in 1% BSA/PBS were then added and incubated for 1 h. Antibodies bound to peptide were detected with anti-Human IgG-HRP secondary antibody diluted in 1% BSA/PBS. Plates were developed with the addition of TMB substrate and stopped by the addition of 2N sulfuric acid. Absorbance at 450 nm was measured on the Synergy Neo2 plate reader (Biotek). A positive signal was identified as an absorbance value above 2-fold of lower limit of detection (defined as average of blank + 3-fold SD of blank).

#### Detection of anti-TREM2 antibody cell binding by flow cytometry

HEK293 overexpressing human TREM2 (HEK293-H6) and HEK293 over expressing GFP were harvested using 0.05% trypsin and incubated at 37°C for 2 h. All cells were centrifuged and washed in FACS buffer (PBS + 0.5% BSA) twice. Mixed cells were resuspended in FACS buffer at a density of 10^6^ cells/ml per cell line. The mixed cell lines were seeded at 100,000 cells per well in a 96-well v-bottom plate and incubated for 20 min at RT. After incubation, the cells were centrifuged and incubated with anti-TREM2 antibodies in a dose titration from 0-300 nM for 45 min on ice. After incubation, cells were centrifuged and washed with FACS buffer three times. The cells were then incubated with secondary antibody (Alexa Fluor 647 AffiniPure F(ab’)2 fragment goat anti-human IgG (H+L), Jackson ImmunoResearch Laboratories, #109-606-088, 1:800 dilution) for 30 min on ice without exposure to light. After incubation, the cells were washed with FACS buffer three times, resuspended in 100 μl of FACS buffer, and analyzed by flow cytometry (BD FACSCanto II, San Jose, CA), for which 50,000 events were obtained for each sample. Mean fluorescence intensity (MFI) per cells were calculated by FLowJO software and used for generation dose response binding curve.

### Antibody treatment

8 h after seeding the cells, they were treated with anti-human TREM2 antibodies (Fig EV1C). Antibodies were diluted in RPMI medium and added to the cells with a final concentration of 20 μg/ml. As control for TREM2 shedding, cells were treated with GM6001 (25 μM, Enzo Life Sciences), or DMSO as a vehicle control. hiMGL were seeded in 6-well plates with 400,000 cells/well. 8 h after seeding, antibodies were diluted in iMGL media and added at a concentration of either 20 or 40 µg/ml. Isotype or TREM2 antibodies were added at a concentration of 40 µg/ml and 24 h after antibody treatment, medium and cells were harvested as previously described.

### Small animal PET/MRI

All rodent PET procedures followed an established standardized protocol for radiochemistry, acquisition times and post-processing (Brendel et al., 2016; Overhoff et al., 2016), which was transferred to a novel PET/MRI system.

All mice were scanned with a 3T Mediso nanoScan PET/MR scanner (Mediso Ltd) with a single-mouse imaging chamber. A 15-min anatomical T1 MR scan was performed at 15 min after [^18^F]-FDG injection or at 45 min after [^18^F]-GE180 injection (head receive coil, matrix size 96 x 96 x 22, voxel size 0.24 x 0.24 x 0.80 mm³, repetition time 677 ms, echo time 28.56 ms, flip angle 90°). PET emission was recorded at 30-60 min p.i. ([^18^F]-FDG) or at 60-90 min p.i. ([^18^F]-GE-180). PET list-mode data within 400-600 keV energy window were reconstructed using a 3D iterative algorithm (Tera-Tomo 3D, Mediso Ltd) with the following parameters: matrix size 55 x 62 x 187 mm³, voxel size 0.3 x 0.3 x 0.3 mm³, 8 iterations, 6 subsets. Decay, random, and attenuation correction were applied. The T1 image was used to create a body-air material map for the attenuation correction. We studied PET images of *Grn* KO mice (n=8), *Trem2* KO mice (n=10), *Double* KO mice (n=10), and WT mice (n=15), all female at an average age of 10.9 ± 1.6 months. Normalization of injected activity was performed by the previously validated myocardium correction method (Deussing et al., 2018) for [^18^F]-GE-180 TSPO-PET and by standardized uptake value (SUV) normalization for [^18^F]-FDG-PET. Groups of *Grn* KO mice, *Trem2* KO and *Double* KO mice were compared against WT mice by calculation of %-differences in each cerebral voxel. Finally, [^18^F]-TSPO-PET and [^18^F]-FDG-PET values deriving from a whole brain VOI (Kleinberger et al., 2017) were extracted and compared between groups of different genotypes by a one-way ANOVA including Tukey post hoc correction.

### CSF collection

Mice were fully anesthetized via an intraperitoneal injection of medetomidine (0.5 mg/kg) + midazolam (5 mg/kg) + fentanyl (0.05 mg/kg). CSF was collected as described previously (Lim et al., 2018). Briefly, subcutaneous tissue and musculature was removed to expose the meninges overlying the cisterna magna. A glass capillary with a trimmed tip (inner diameter is approximately 0.75 mm) was used to puncture the membrane, and CSF was allowed to flow into the capillary for approximately 10 min. After collection, CSF was centrifuged at 1000 g for 10 min, assessed macroscopically for blood contamination, aliquoted (5 μl) in propylene tubes, snap-frozen in liquid nitrogen, and stored at −80°C until use.

#### CSF neurofilament light chain analysis

NfL levels were quantitatively determined in CSF samples using the Simoa NF-light Advantage Kit (Quanterix, #103186) following the manufacturer’s instructions. CSF samples were diluted 1:10 in sample dilution buffer and mixed with Simoa detector reagent and bead reagent, following an incubation at 30°C for 30 min, shaking at 800 rpm. Plates were washed with Simoa washing buffer A and SBG reagent from the kit was added. Following a 10 min incubation at 30°C, shaking at 800 rpm, plates were washed twice and sample beads were resuspended in Simoa wash buffer B. NfL concentrations were measured after a 10 min drying at RT using the Simoa DH-1 analyzer (Quanterix).

### Gene expression profiling of total brain

Adult mice were perfused transcardially with PBS and dissected brains were snap frozen in liquid nitrogen. Snap frozen brains were mechanically powdered in liquid nitrogen. Total RNA was isolated using the RNeasy Mini Kit (Qiagen, #74104) and 60 ng of total RNA per sample were subjected to gene expression profiling using the nCounter® Neuropathology panel from NanoString (NanoString Technologies). Gene expression levels in each sample were normalized against the geometric mean of four housekeeping genes including *Asb10, Cltc, Hprt1* and *Tubb5* using the *nSolver Analysis Software*, version 4.0. *Gusb* was excluded because of significant changes in *Grn* KO and *Double* KO mice.

### Gene expression profiling of primary microglia

CD11b+ and FCRL+ primary microglia were isolated from adult mouse brain. Mice were perfused transcardially and brains were collected into ice-cold HBSS (ThermoFisher, #14175095). Brain tissue was mechanically dissociated into single-cell suspension using Potter-Elvehjem homogenizers with PTFE Pestle Microglia cell pellets were resuspended 70 % Percoll and overlayed with equal volumes of 40 % Percoll. Microglia were enriched at the interface of 70% (v/v) to 40% (v/v) Percoll after centrifugation (at 18°C, 300 x g for 30 min; slow acceleration and deceleration:3) (Mazaheri et al., 2017). Microglia were collected, filtered through 100 μm cell strainers and washed with blocking buffer (0.2% BSA in HBSS). Cells were then consecutively stained with FCRLs monoclonal rat antibody (Butovsky et al., 2014) (30 min), goat anti-rat APC antibody (Biogegend, # 405407) (20 min) and Cd11b PeCy7 antibody (BD, #553142) (20 min) on ice. Cells were then washed and resuspended in 0.5 ml blocking buffer and subjected to cell sorting. Sorted CD11b+ and FCRL+ cells were pelleted by centrifugation and snap frozen in liquid nitrogen, stored at −80°C until further use. Following total cell lysis in 1:3 diluted RLT buffer (Quiagen, RNeasy Mini Kit), 10.000 cells in 4 μl volume were subjected to gene expression profiling with the nCounter® customized panel from NanoString (NanoString Technologies). We generated an nCounter panel for analyzing gene expression of 65 microglial activation related genes including five (*Asb10, Cltc, Hprt1 Tubb5* and *Gusb*) housekeeping genes. Gene expression levels in each sample were normalized against the geometric mean of four housekeeping genes using the *nSolver Analysis Software*, version 4.0. *Gusb* was excluded because of significant changes in *Grn* KO and *Double* KO mice.

### Gene expression profiling of human hiMGL

hiMGL were seeded into 12-well plates and incubated for 3 h at 37°C (5% CO_2_). Thereafter, microglia were treated with TREM2antagonistic antibodies and isotype control for 24 h. After treatment, cells were collected and RNA was isolated using the E.Z.N.A HP Total RNA Kit (Omega Bio-tek) according to the manufacturer’s instructions. Following isolation, RNA quality was determined using a 4200 TapeStation (Agilent) and gene expression profiling with the nCounter® customized panel from NanoString (NanoString Technologies) was performed. We generated an nCounter panel for analyzing gene expression of 82 microglial related genes and 8 housekeeping genes. Gene expression levels in each sample were normalized against the geometric mean of five housekeeping genes including *CLTC, HPRT1, RPL13A, TBP* and *PPIA* using the *nSolver Analysis Software*, version 4.0. *CALR, TUBB5* and *YWAHZ* were excluded because of significant changes in *Grn* KO and WT microglia.

### Lipid analysis by liquid chromatography-mass spectrometry (LCMS)

#### Sample preparation for LCMS

For LCMS sample preparation, 10 mg of brain powder prepared from whole brain powered homogenates was mixed with 400 ml of methanol spiked with internal standards and homogenized with a 3 mm tungsten carbide bead (shaken at 25 Hz for 30 seconds). The methanol fraction was then isolated via centrifugation (20 min at 4°C, 14,000 x g, followed by transfer of supernatant to a 96 well plate, 1 h incubation at −20°C followed by an additional 20 min centrifugation (4,000 x g at 4°C) and transferred to glass vials for LCMS analysis. For analysis of a GlcCer/GalCer panel, an aliquot of the methanol fraction was dried under N_2_ gas and then resuspended in 100 ml of 92.5/5/2.5 CAN/IPA/H2) (MS grade) with 5 mM ammonium formate (MS grade) and 0.5% formic acid (MS grade).

Unless otherwise noted, relative quantification of lipids and metabolites were performed using the Shimadzu Nexera X2 LC system (Shimadzu Scientific Instrument) coupled to Sciex QTRAP 6500+ mass spectrometer (Sciex).

#### Lipidomic analysis

For each analysis, 5 µl of sample was injected on a BEH C18 1.7 µm, 2.1×100 mm column (Waters Corporation) using a flow rate of 0.25 ml/min at 55°C. Mobile phase A consisted of 60:40 acetonitrile/water (v/v); mobile phase B consisted of 90:10 isopropyl alcohol/acetonitrile (v/v). These buffers were fortified with 10 mM ammonium formate with 0.1% formic acid (positive ionization) or with 10 mM ammonium acetate (negative ionization). The gradient was programmed as follows: 0.0–8.0 min from 45% B to 99% B, 8.0–9.0 min at 99% B, 9.0–9.1 min to 45% B, and 9.1–10.0 min at 45% B. Source settings were as follows: curtain gas at 30 psi; collision gas was set at medium; ion spray voltage at 5500 V (positive mode) or 4500 V (negative mode); temperature at 250°C (positive mode) or 600°C (negative mode); ion source gas 1 at 55 psi; ion source gas 2 at 60 psi. Data acquisition was performed using Analyst 1.6.3 (Sciex) in multiple reaction monitoring mode (MRM). Area ratios of endogenous metabolites and surrogate internal standards were quantified using MultiQuant 3.02 (Sciex).

### Protein analysis and Western blotting

Cell pellets obtained from human primary monocytes, cultured hiMGL or aliquots of powdered frozen brains were lysed in Triton lysis buffer (150 mM NaCl, 50 mM Tris-HCL, pH 7.6, 2 mM EDTA, 1 % Triton X-100) supplemented with protease inhibitor (Sigma-Aldrich). Lysates were incubated on ice for 30 min and then centrifuged at 17,000 x g for 15 min at 4°C. For sequential biochemical protein extraction of soluble, less-soluble and insoluble proteins, brain powder was lysed in high salt (HS) buffer (0.5 M NaCl, 10 mM Tris-HCL pH 7.5, 5 mM EDTA, 1 mM DTT, 10% sucrose), then RIPA buffer (150 mM NaCl, 20 mM Tris-HCL pH7.4, 1% NP-40, 0.05% Triton X-100, 0.5% sodium-desoxycholate, 2.5 mM EDTA) followed by urea buffer (30 mM Tris-HCL pH 8.5, 7 M urea, 2 M thiourea) as described previously (Gotzl et al., 2014). For membrane preparation of hiMG pellets were resuspended with hypotonic buffer (10 mM Tris, pH 7.4, 1 mM EDTA, pH 8.0, 1 mM EGTA, pH 8.0) and incubated on ice for 30 min, vortexed every 10 min, followed by a freeze-thaw cycle and centrifuged at 17,000 x g for 45 min at 4^°^C. The supernatants were collected (cytosolic fraction) and the pellet (membrane fraction) resuspended in STEN lysis buffer and incubated on ice for 20 min. Insoluble proteins were pelleted at 17,000 x g for 20 min at 4^°^C and the supernatant (membrane fraction) was collected and used for further analysis. Protein concentrations were determined using the BCA protein assay (Pierce, ThermoFisher). Equal amounts of protein adjusted to lysis buffer were mixed with Laemmli sample buffer supplemented with β-mercaptoethanol. Proteins were separated by SDS-PAGE and transferred onto polyvinylidene difluoride membranes (Immobilon-P, Merck Millipore). Proteins of interest were detected using the following primary antibodies: goat anti-TREM2 (R&D Systems, Inc., #AF1828), rabbit anti-PGRN (ThermoFisher, #40-3400), goat anti-CatD (R&D, #AF1029), mouse anti-βActin (Sigma, #A5316) and rabbit anti-Calnexin (Stressgene, #SPA-860) followed by incubation with horseradish peroxidase-conjugated secondary antibodies and ECL Plus substrate (ThermoFisher, Pierce ECL Plus Western Blotting Substrates). For quantification, images were taken with a Luminescent Image Analyzer LAS-4000 (Fujifilm Life Science, Tokyo, Japan) and evaluated with the Multi GaugeV3.0 software (Fujifilm Life Science, Tokyo, Japan).

### ELISA-based quantification of sTREM2 and PGRN

sTREM2 in conditioned media was quantitated using the Meso Scale Discovery Platform as described previously (Schlepckow et al., 2020). Briefly, streptavidin-coated small spot 96-well plates were blocked overnight at 4°C, incubated with 0.125 μg/ml biotinylated polyclonal goat anti-human TREM2 capture antibody (R&D, #BAF1828). After washing, plates were incubated with samples and standards for 2 h at RT. If cells were antibody treated, samples and standards were previously mixed 9:1 with denaturing buffer (200 mM Tris-HCL pH 6.8, 4% (w/v) SDS, 40% (v/v) glycerol, 2% (v/v) β-mercaptoethanol, 50 mM EDTA) and boiled at 95°C for 5 min to dissociate and denature TREM2 antibodies bound antibodies. Plates were washed before incubation for 1 h at RT with 1 μg/ml mouse anti-human TREM2 antibody (SantaCruz Biotechnology, B-3 SCBT-373828). After washing, plates were incubated with a SULFO-TAG-labeled anti-mouse secondary antibody (MesoScaleDiscovery, R32AC-5) for 1 h at RT. After additional washing steps, 1x Meso Scale Discovery Read buffer was added and the light emission at 620 nm after electrochemical stimulation as measured with an Meso Scale Discovery Sector Imager 2400 reader.

PGRN levels were determined using a previously described protocol (Gotzl et al., 2019) using the following antibodies: a biotinylated polyclonal goat anti-human PGRN antibody (R&D, #BAF2420) at 0.2 μg/ml as capture antibody, a mouse anti-human PGRN antibody (R&D, #MAB2420) as detection antibody and a SULFO-TAG-labeled anti-mouse (MesoScaleDiscovery, #R32AC-5) as secondary antibody.

### p-Syk AlphaLISA

Phosphorylated SYK (p-Syk) was measured using the AlphaLISA SureFire Ultra p-Syk Assay Kit (PerkinElmer, #ALSU-PSYK-A-HV) following the manufacturer’s instructions. Briefly, differentiated human macrophages were plated in 100 μl media at a density of 50,000 cells/well. hiMGL were plated in iMGL media at a density of 30,000 cells/well, in 96-well plates and incubated overnight at 37°C in a cell culture incubator. Plates were then washed three times with HBSS and 50 μl of liposome (1 mg/ml)/antibody (20 μg/ml) mix was added to the cells. Following an incubation at 37°C for 1 h for macrophages, or 5 min for hiMGL, treatment solutions were removed and cells were lysed with 40 μl lysis buffer supplemented with protease inhibitor mix (Sigma) and phosphatase inhibitor (PhosSTOP, Roche) for 30 min at 4°C. Equal volumes of lysate were then subjected to analysis using an EnSpire Multimode Plate Reader (PerkinElmer).

### Liposome Preparation

POPC/POPS (7:3) liposomes at 10 mg/ml were prepared as follows: 7 mg POPC and 3 mg POPS were dissolved in chloroform followed by thorough evaporation of solvent. The lipid mixture was then resuspended in 1 ml HBSS, and suspensions were extruded using 100 nm polycarbonate membranes (Whatman, #800309) and a LiposoFast extruder device (Sigma-Aldrich) to generate large unilamellar vesicles.

### Cathepsin activity assay

hiMGL cell pellets or powdered mouse brain tissue was used for cathepsin D fluorescence-based activity assays (Abnova) as described previously (Gotzl et al., 2018). Mouse brain tissue was homogenized using precellys lysing kit (Bertin Instruments, #P000933-LYSK0-A).

### GCase activity assay

Brain powder was lysed in GCase lysis buffer (150 mM NaCl, 20 mM Tris-HCl (pH 7.5), 1% Triton X-100, 1 mM EDTA, 1 mM EGTA) and protein concentrations were determined using the BCA assay. Lysates were adjusted to 2 mg/ml. Lysates were diluted 12,5-fold in GCase activity buffer (100 mM Phosphate Citrate buffer pH 5.2, 0.5% Sodium Taurocholate, 0.25% Triton-X 100) and 4-Methylumbelliferyl β-D-glucopyranoside stock solution (30 mM; Sigma-Aldrich, M3633-1G, stock solution in DMF) was diluted 3-fold in the GCase activity buffer. 90 μl of the diluted lysates and 10 μl of the diluted 4-Methylumbelliferyl β-D-glucopyranoside were added to a 96-well plate. Plates were incubated for 15 min at 37 °C. Signal intensities were measured continuously for 1 h (Ex 365 nm/Em 455 nm).

### Phagocytosis assays

Microglial phagocytosis was determined using the IncuCyte S3 Live-Cell Analysis System (Sartorius). hiMGL cells were plated in 96-well plates at 30% confluency. Cells were incubated at 37°C and the confluency (pre-treatment) of each well was determined with the IncuCyte S3 3 h after seeding. Thereafter, TREM2 antagonistic antibodies and isotype control were added at 20 and 40 µg/ml. 18 h after antibody treatment pHrodo labeled myelin (5 µg/ml) was added to the hiMGL cells and images of fluorescence and phase were captured at 4X in the IncuCyte S3 live cell imager every 15 min. Using IncuCyte 2020B software (Satorius), image masks for phase and fluorescent signal (phagocytosis of pHrodo-labelled myelin) were acquired, and the fluorescent signal was normalized to cell confluency (cell body area), which was measured before the antibody treatment.

### pHrodo labeling of myelin

Myelin was labeled with amine reactive pHrodo^TM^ Red succinimidyl ester (ThermoFisher) for 45 min at RT (protected from light). Labeled myelin was washed with PBS and either directly used or stored in aliquots at −80°C.

### Immunohistochemistry and image acquisition

Mice were transcardially perfused with PBS and brains were dissected into two hemispheres. One hemisphere was snap frozen and stored at −80°C until further use. The other hemisphere was immersion-fixed for 24 h in 4% paraformaldehyde, washed with PBS and incubated 30% sucrose for 48 h for cryoprotection. After freezing, brains were cut into 50 μm or 100 μm coronal sections using a vibratome (Leica Biosystems), collected in PBS and stored at 4°C until further use. For visualizing lipofuscin, 50 μm free-floating sections were incubated with 5% donkey serum in PBS overnight at 4°C with slow agitation. After washing, tissue sections were stained with DAPI for 10 min at RT and mounted onto slides using ProlongTM Gold Antifade reagent (ThermoFisher, #P36961). For morphological analysis of microglia, 100 μm sections were blocked with goat serum blocking buffer (2% goat serum, 0.05% Tween 20 in 0.01 M PBS pH 7.2-7.4) and stained with anti-Iba1 in primary antibody buffer (1% bovine serum albumin, 0.1% gelatin from cold water fish skin, 0.5% Triton X-100 in 0.01 M PBS pH 7.2-7.4) and anti-rabbit IgG coupled Alexa Flour 555 in secondary antibody buffer (0.05% Tween 20 in 0.01 M PBS pH 7.2-7.4). After washing, tissue sections were stained with DAPI for 10 min at RT and mounted onto slides using ProlongTM Gold Antifade reagent (Firma). Images were acquired using a LSM800 Zeiss confocal microscope and the ZEN 2011 software package (blue edition). For lipofuscin analysis, five images were taken per slide using a 20x objective at 2048 x 2048 pixel resolution. Total fluorescence was quantified using the FIJI software (ImageJ).

### Automated analysis of microglia morphology

For morphological analysis of microglia, three z-stack images per animal (n=3) were recorded with a 40x objective in a resolution of 1024 x 1024 pixels (x-y-pixel size = 0.15598 µm) and a slice distance (z) of 0.4 µm. The raw confocal z-stacks were then analyzed using the Microglia Morphology Quantification Tool (MMQT) for automated analysis of microglial morphology as previously described (Heindl et al., 2018). The algorithm was run in MATLAB (Version R2016b). To identify the most discriminating features, a receiver operating characteristic (ROC) analysis was performed in R (version 4.0.3) for calculating the area under the curve (auc) between the groups “WT” and “*Double* KO“. Statistical analysis of group difference for the morphological scores “Branch volume” (auc = 0.72), “Sphericity score” (auc = 0.82), “Branch length” (auc = 0.69) and “Number of branch nodes” (auc = 0.80) was performed using the Wilcoxon rank sum test with continuity correction and Bonferroni post-hoc correction for multiple testing in R (version 4.0.3).

### Statistical analysis

Data were analyzed using GraphPad Prism 9. If no other test of significance is indicated, for statistical analysis of two groups of samples the unpaired, two-tailed student’s t-test was performed. For comparison of more than two groups, one-way ANOVA and Dunnett’s or Tukey’s post hoc test was used. Statistical significance was set at *, *p* < 0.05; **, *p* < 0.01; and ***, *p* < 0.001; and ****, *p* < 0.0001.

## Acknowledgement

This work was supported by grants from the Deutsche Forschungsgemeinschaft (DFG, German Research Foundation) under Germany’s Excellence Strategy within the framework of the Munich Cluster for Systems Neurology (EXC 2145 SyNergy – ID 390857198) (to CH and DP), a Koselleck Project HA1737/16-1 (to CH), BR4580/1-1 (to MB), the Helmholtz-Gemeinschaft Zukunftsthema ‘Immunology and Inflammation’ (ZT-0027) (to CH), Alzheimer’s Association (to CH and DP), Vascular Dementia Research Foundation (to DP), and the donors of the ADR AD2019604S, a program of the BrightFocus Foundation (to DP). AR is supported by a Ph.D. stipend from the Hans and Ilse Breuer Foundation. MB was supported by the Alzheimer Forschung Initiative e.V (grant number 19063p). The authors like to thank Michael Heide and Oliver Weigert (Core Facility “Digital Single Molecule/NanoString Technologies; Deutsches Konsortium für Translationale Krebsforschung, Partner Site München, Labor für Experimentelle Leukämie- und Lymphom-Forschung (ELLF)) for supporting NanoString measurements, Ludovico Cantuti-Castelvetri and Mikael Simons for preparing labeled myelin, Jane Hettinger and Alba Simats for performing CSF withdrawal. LC/MS-based lipidomics analyses was supported by Sonnet Davis. Antibody discovery, material generation, and characterization of TREM2 antagonist antibodies were supported by Josh Park, Do Jin Kim, Yaneth Robles, Rachel Prorok, Steve Lianoglou, Cathal Mahon, and Tina Giese from Denali Therapeutics.

## Author contributions

CH, AC and AR conceived the study and analyzed the results. CH wrote the manuscript with further input from AR, AC, GDP, KMM, JWL, SR and DP. AR performed and analyzed Western Blots, ELISAs, enzyme activity assays, mRNA isolation, NanoString experiments and immunofluorescence on all mouse samples. AR isolated and performed all experiments of human derived macrophages and analyzed hiMGL NanoString data. With supervision of DP, SR generated and validated *GRN* KO hiPSC, differentiated into hiMGL and performed and analyzed Western Blots, ELISAs and enzyme activity assays. JK and GK helped to establish hiMGL cell differentiation. KMM and BVL conducted, generated and validated antagonistic TREM2 antibodies. TL and JS performed lipidomic analysis. MAV and SH performed NanoString and phagocytosis assays on hiMGL cells. JG, KW, AZ, and MB conducted, performed and analyzed PET imaging. SH performed automated analysis on microglia morphology. JL, KB, JDS, EW and LR identified FTLD patients and performed sequencing analysis. BN performed NfL measurements.

## Expanded View Figure Legends

**Figure EV1.**
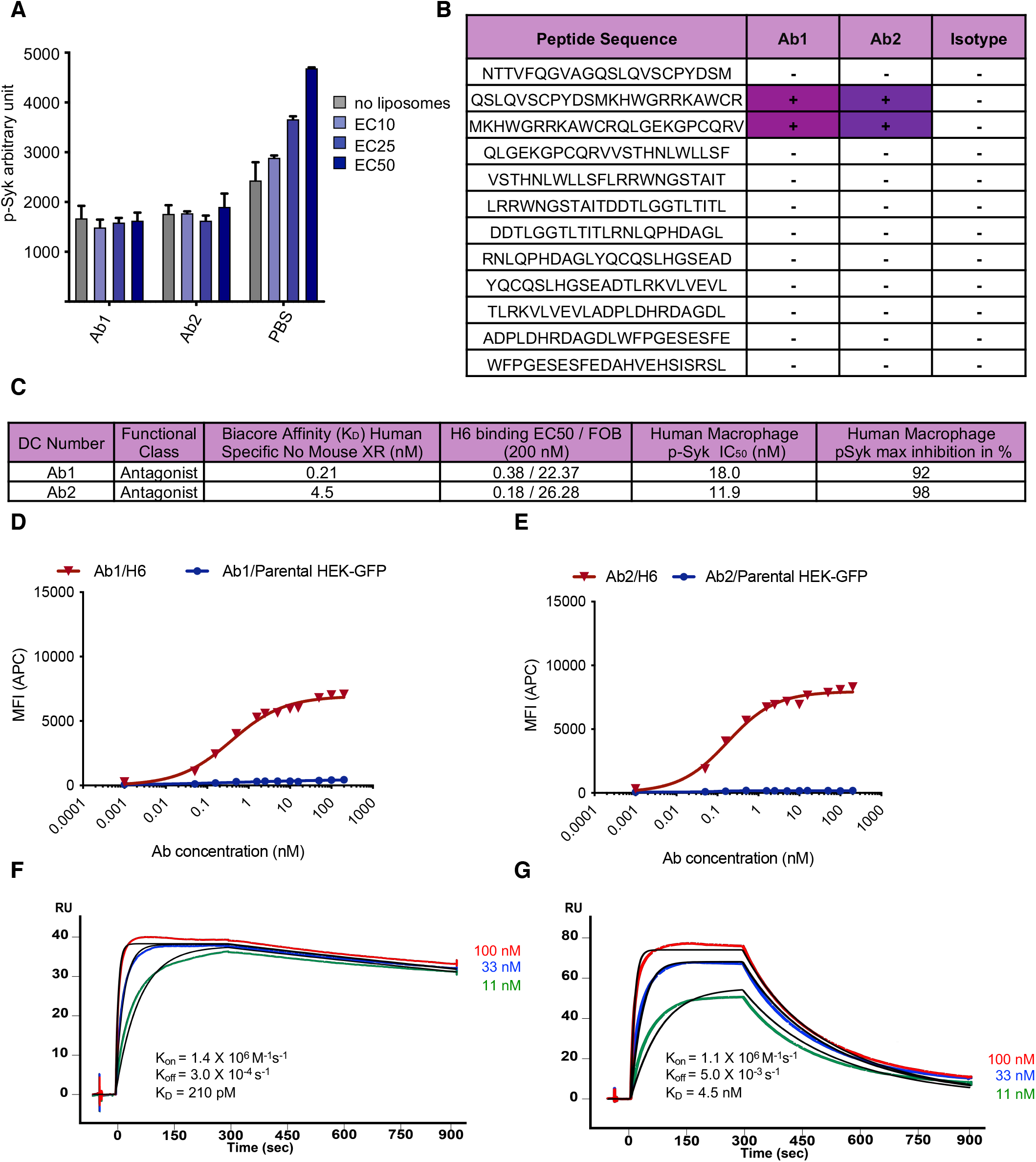
Generation and characterization of TREM2 antagonistic antibodies. **A.** AlphaLISA quantification of p-Syk levels in HEK293 cells over-expressing TREM2-DAP12 upon treatment with Ab1 or Ab2 with three doses of liposomes (EC10: 0.0189527 mg/ml, EC25: 0.0633607 mg/ml or EC50: 0.211731). PBS was used as a negative control. Data represent the mean ± SEM (n=2). **B.** Biochemical binding data of TREM2 IgV peptides to antagonistic antibodies Ab1 and Ab2, as well as isotype control. Positive data represent binding level above a threshold of 2*LLOD (lower limit of detection), from an average of 3 independent experiments. **C.** Table of Ab1 and Ab2 with Biacore binding affinities, cell binding affinities, liposome signaling (pSyk) inhibition (EC50 and maximal inhibition) in human macrophage (n=3 independent experiments). **D., E.** Cell binding dose-response curves generated by FACS in HEK293 cells over-expressing TREM2/DAP12 vs parental lines shown as mean fluorescence intensity (MFI). **F., G.** Biacore binding measurements of immobilized antagonist antibody to 3 concentrations of recombinant hTREM2-ECD

**Figure EV2.**
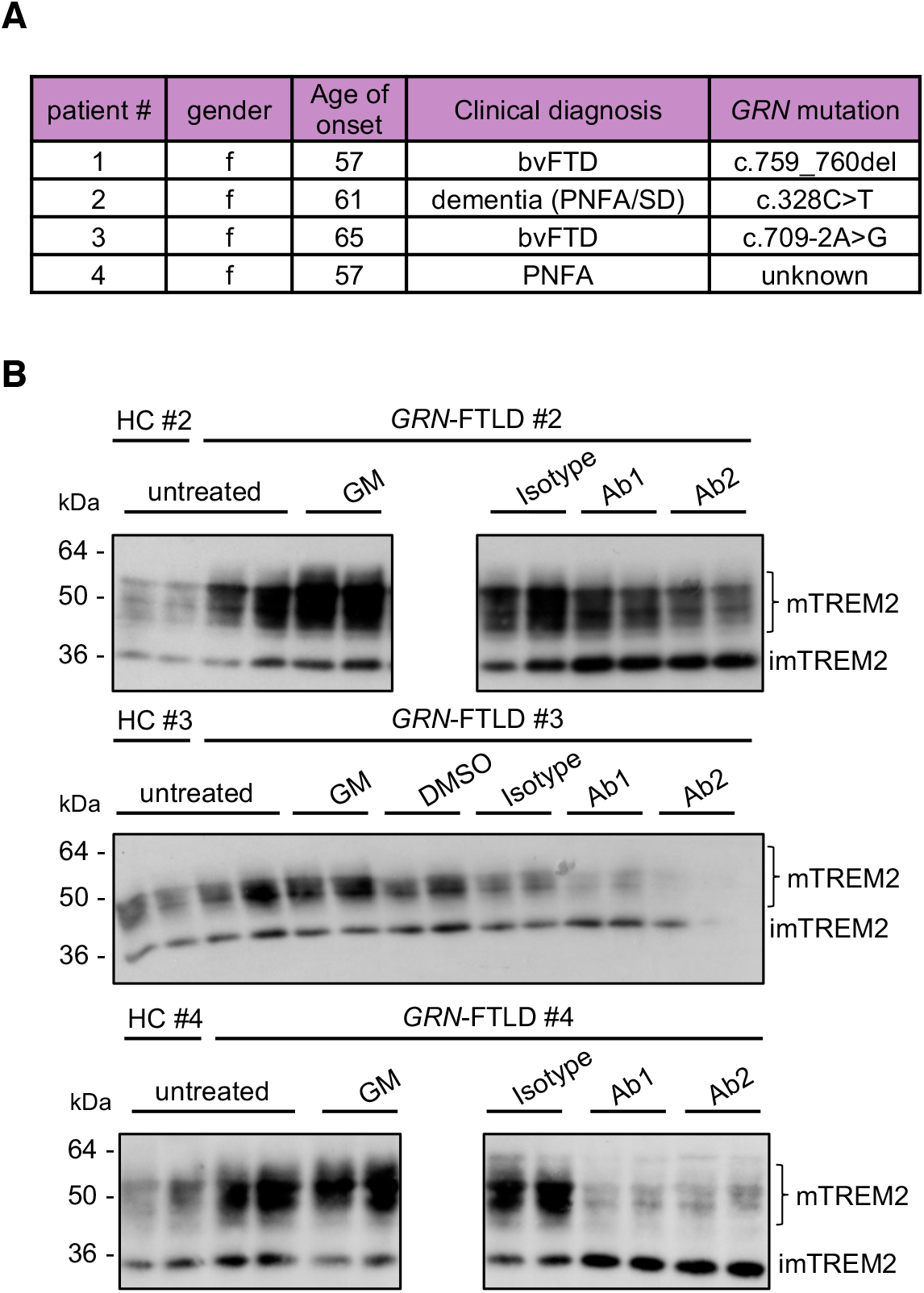
*GRN*-FTLD patients. **A.** Clinical data and mutation status of the four identified GRN mutation carriers. The mutation status of patient #4 is unknown, nevertheless, PGRN serum levels are significantly reduced compared to healthy controls (Fig. 3C). **B.** Western blot analysis of TREM2 using lysates of cultured human macrophages isolated from *GRN* mutation carriers (patients #2 to #4) (A) and healthy control upon treatment with Ab1 and Ab2. An isotype antibody was used as a negative control. ADAM protease inhibition (GM) does not further increase mTREM2 levels in PGRN mutation carriers. Equal amounts of protein were loaded.

**Figure EV3.**
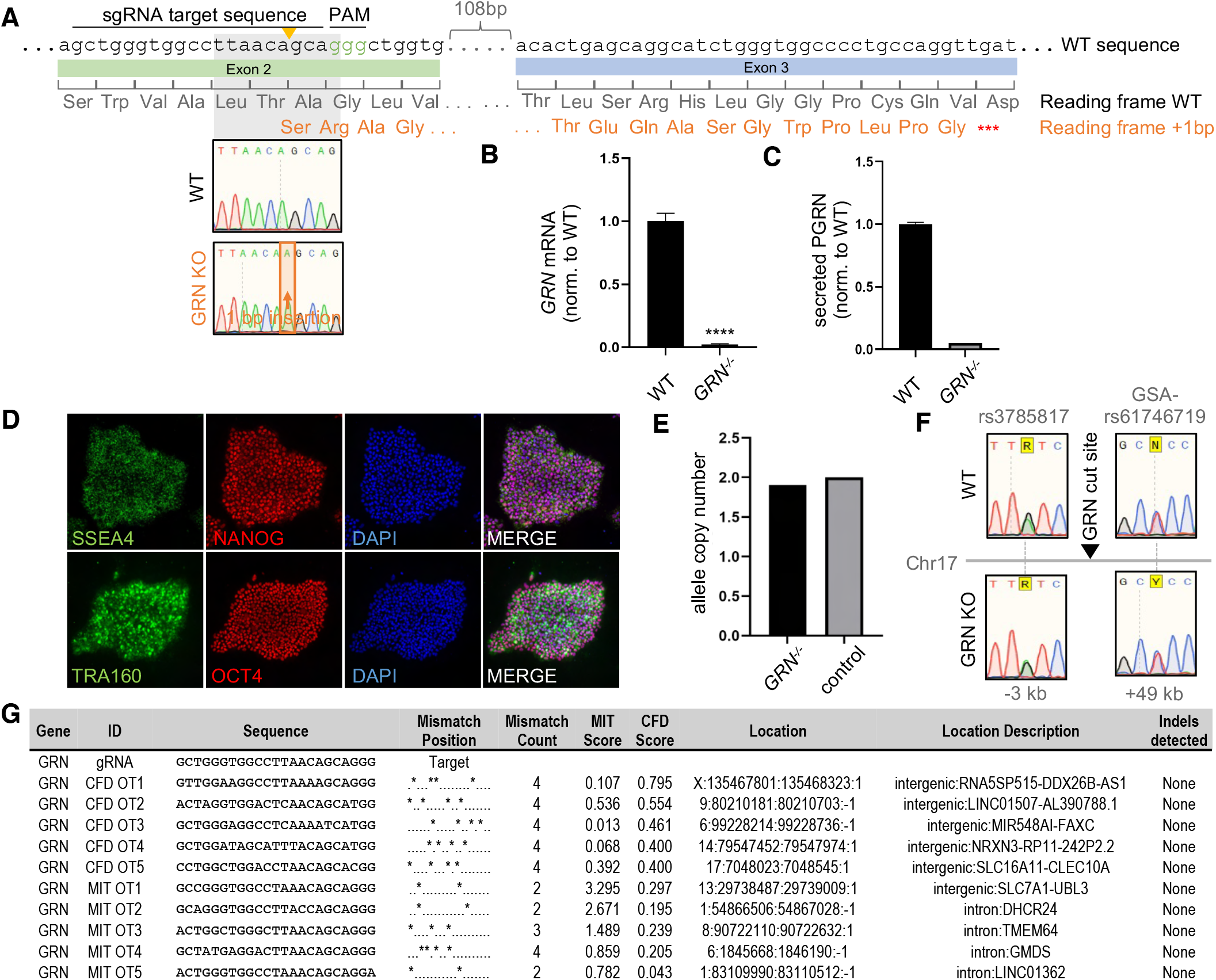
Generation and characterization of *GRN*^-/-^ iPSC line. **A.** *GRN* knockout generation strategy: *GRN* was targeted in exon 2 by a sgRNA (target and PAM sequence indicated), leading to a one base pair insertion in the *GRN*^-/-^ line. The resulting frameshift exposes a nearby stop codon. **B.** *GRN* mRNA transcipt levels in WT and *GRN^-/-^* hiMGL normalized to WT, as measured by qPCR (n=3). **C.** ELISA-mediated quantification of secreted PGRN in conditioned media of WT and *GRN^-/-^* hiMGL (n = 3). **D.** Immunofluorescence analysis of pluripotency markers SSEA4, NANOG, TRA160, and OCT 4 with DAPI in *GRN*^-/-^ iPSCs. **E., F.** Analysis of CRISPR-mediated on-target effects by qgPCR quantitation of allele copy number (E) and Sanger sequencing of SNPs near the edited locus in WT and *GRN^-/-^* iPSC lines (F) shows maintenance of both alleles after editing. **G.** List of top five most similar off-target sites ranked by the CFD and MIT prediction scores, respectively. No off-target editing was detected by Sanger sequencing. Data represent the mean ± SD. For statistical analysis the unpaired, two-tailed student’s t-test was performed. Statistical significance was set at *, *p* < 0.05; **, *p* < 0.01; ***, *p* < 0.001; ****, *p* < 0.0001, and ns, not significant.

**Figure EV4.**
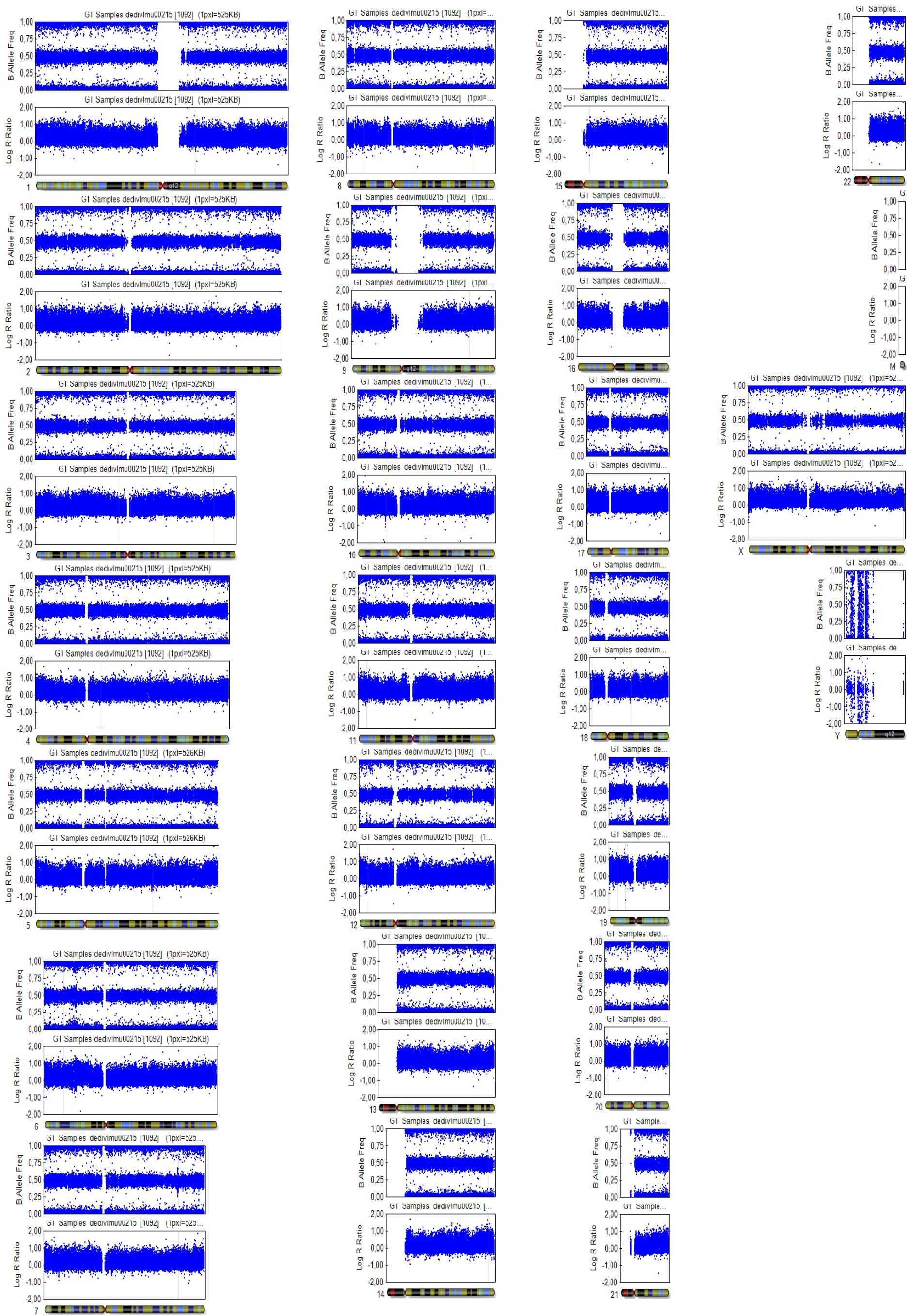
Molecular karyotyping of *GRN*^-/-^ iPSC line. B allele frequencies (BAF) and Log R ratios are shown for each chromosome in the *GRN*^-/-^ iPSC line. Measured SNPs are indicated by blue dots. BAF values indicate normal zygosities on all chromosomes and Log R ratios confirm absence of detectable deletions or insertions. Overall, the karyotype shows no chromosomal aberrations.

**Figure EV5.**
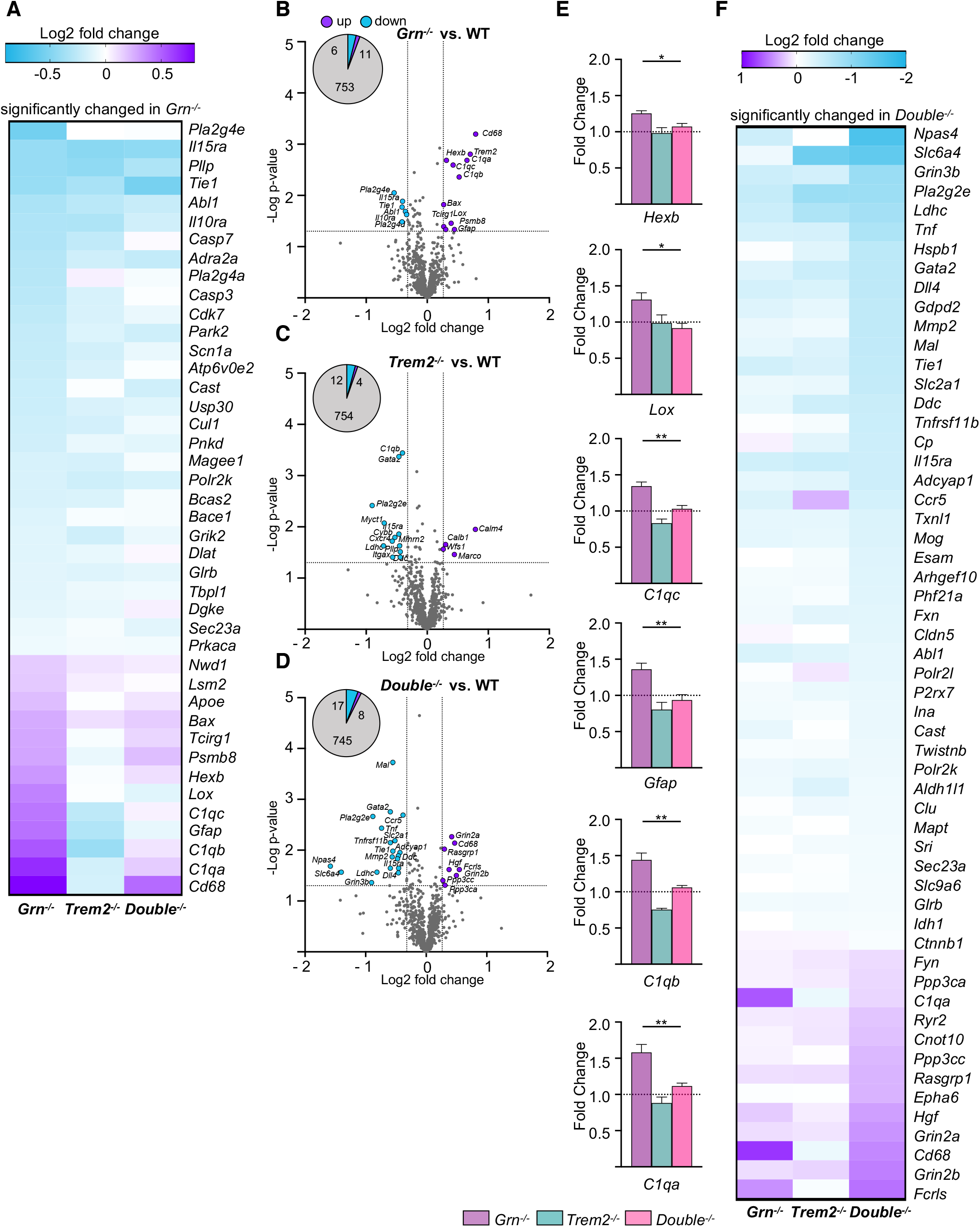
Differential gene expression in *Grn*^-/-^ and *Trem2*^-/-^ is partially rescued in *Double* KO mice. **A.** Heatmap of significantly changed gene transcripts in whole brain of 6-month-old *Grn^-/-^, Trem2^-/-^* and *Double^-/-^* mice in comparison to WT (n=6 per genotype) detected using the Neuropathology panel by NanoString. Only genes of significantly changed transcript levels in *Grn^-/-^* in comparison with WT mice are displayed. Changes in expression were only considered if changes are above 20%. mRNA counts for each gene and sample were normalized to the mean value of WT followed by a log2 transformation. **B.** Volcano blot presentation of the differently expressed transcripts in brains of 6-month-old *Grn^-/-^* mice compared to WT brain mRNA (n=4 per genotype). 17 out of 752 analyzed genes are significantly changed more than 20%, with 11 genes upregulated (purple) and 6 genes downregulated (blue). **C.** Volcano blot presentation of the differently expressed genes in brain mRNA of 6-month-old *Trem2^-/-^* mice in comparison to WT brain mRNA (n=4 per genotype). 16 out of 752 analyzed genes are significantly changed by more than 20%, with 4 genes upregulated (purple) and 12 genes downregulated (blue). **D.** Volcano blot presentation of the differently expressed genes in brain mRNA of 6-month-old *Double^-/-^* mice in comparison to WT brain mRNA (n=4 per genotype). 25 out of 752 analyzed genes are significantly changed by more than 20%, with 8 genes upregulated (purple) and 17 genes downregulated (blue). **E.** Transcript levels of selected significantly rescued genes in *Double^-/-^* vs. *Grn^-/-^* brain mRNA. Transcript expression is normalized to the mean of the WT cohort. **F.** Heatmap of significantly changed transcripts in whole brain mRNA of 6-month-old *Grn^-/-^, Trem2^-/-^* and *Double^-/-^* mice in comparison to WT (n=6 per genotype) detected by the Neuropathology panel by NanoString. Only genes significantly changed in *Double^-/-^* in comparison to WT mice are displayed. Changes in expression were only considered if changes are above 20%. RNA counts for each gene and sample were normalized to the mean value of WT followed by a log2 transformation. Data represent mean ± SEM. For statistical analysis in **B** - **D** the unpaired, two-tailed student’s t-test was performed, and in **E** one-way ANOVA with Dunnett’s post hoc test was used to compare *Grn^-/-^, Trem2^-/-^* and *Double^-/-^* whole brain mRNA expression. Statistical significance was set at *, *p* < 0.05; **, *p* < 0.01; ***, *p* < 0.001; ****, *p* < 0.0001, and ns, not significant.

